# MAKR6 integrates TMK and CAMEL/CANAR signalling for auxin canalization in Arabidopsis

**DOI:** 10.1101/2025.10.07.680881

**Authors:** Zengxiang Ge, Lilla Koczka, Ewa Mazur, Gergely Molnar, Dmitrii Vladimirtsev, Nada Kassem, Sara Ait Ikene, Lukáš Fiedler, Jiří Friml

**Affiliations:** Institute of Science and Technology Austria (ISTA), Klosterneuburg, 3400, Austria; Mendel Centre for Plant Genomics and Proteomics, Central European Institute of Technology (CEITEC), Masaryk University, Kamenice 5, CZ-625 00 Brno, Czech Republic; National Centre for Biomolecular Research, Faculty of Science, Masaryk University, Kamenice 5, CZ-625 00 Brno, Czech Republic; Institute of Biology, Biotechnology and Environmental Protection, Faculty of Natural Sciences, University of Silesia in Katowice, Katowice, 40-032, Poland

**Keywords:** auxin canalization, auxin transport, PIN polarity, MAKR6, receptor-like kinase

## Abstract

Adaptive plant development is orchestrated, among others, by directional, intercellular transport of the phytohormone auxin. Self-organizing development, such as flexible vasculature formation, depends on so-called auxin canalization, manifested by the gradual formation of auxin transport channels through feedback between auxin signalling and transport. Herein, we identify MAKR6 as an important, novel component in this feedback. *MAKR6* expression accumulates strongly in vascular cells and is tightly regulated by auxin via the Aux/IAA-ARF-WRKY23 transcriptional network. MAKR6 is required for auxin canalization-dependent processes, including leaf venation, vasculature regeneration, and *de nov*o auxin channel formation from local auxin sources. Mechanistically, MAKR6 interacts with the PIN1 auxin transporter, modulating its trafficking and polarization. MAKR6 also associates with and integrates two key receptor-like kinase complexes involved in canalization, TMK1/4 and the CAMEL-CANAR. Together, our study establishes MAKR6 as a multifaceted regulator that couples transcriptional auxin signalling to PIN1 repolarization and coordinates multiple RLK-mediated signalling pathways during canalization. This provides mechanistic insights into auxin canalization and exemplifies a framework for exploring similar regulatory nodes in other developmental contexts.

## Main

The auxin canalization hypothesis explains self-reinforcing processes, including vasculature formation and regeneration, and possibly also embryogenesis and organogenesis^1–3^. In this model, cells adjust the polarity of PIN auxin transporters to direct auxin flow away from auxin sources, gradually forming auxin channels, thereby providing positional cues for vasculature formation^4–8^.

Despite its importance and conceptual uniqueness, the molecular mechanism of auxin canalization remains poorly understood. It integrates TIR1/AFB-Aux/IAA-ARF transcriptional auxin signalling with polar auxin transport by PIN proteins^5,9,10^. For example, the transcription factor WRKY23 acts downstream to regulate PIN repolarization^11^. One WRKY23 target encodes CAMEL (Canalization-related Auxin-regulated Malectin-type RLK), a receptor-like kinase (RLK) mediating canalization-dependent processes, including vasculature regeneration and vein patterning. CAMEL interacts with CANAR (Canalization-related Receptor-like kinase), regulating PIN1 trafficking and polarization^12^. Cell-surface auxin perception by ABP1 (Auxin-Binding Protein 1) and its partners, TMKs (TransMembrane Kinases), is also required for the formation of auxin channels and vasculature^13,14^. TOW (Target Of WRKY23) is an essential component of canalization, promoting PIN interaction with CAMEL and TMK to stabilize PINs at the cell surface^15^. Identification of all these components underscores the mechanistic complexity, yet the specific cues beyond auxin, additional players, and mutual regulation remain unclear.

Intrinsically disordered BKI1 (BRI1 Kinase Inhibitor1) and MAKR (Membrane-Associated Kinase Regulator) proteins modulate the activity of PM-localized RLKs in diverse processes^16^. For instance, BKI1 regulates brassinosteroids (BRs) and EPFs (Epidermal Patterning Factors) signalling, while MAKR2/4/5 act in the gravity-induced root bending, lateral root initiation, and protophloem differentiation, respectively^17–20^. Although RLKs such as CAMEL, CANAR, and TMKs have provided key molecular insights into auxin canalization, their regulators, such as MAKRs, are poorly defined.

In this study, we identified MAKR6, a downstream component of transcriptional auxin signalling, as an important regulator of canalization-dependent processes, including venation and vascular regeneration. MAKR6 functions as a molecular hub, integrating TMK1/4 and CAMEL-CANAR signalling complexes and controlling PIN1 trafficking and polarity. By bridging auxin signalling to PIN polarization and coordinating mutual RLK functions, MAKR6 reinforces the feedback loop required for auxin canalization, ultimately facilitating directional auxin flow and proper vascular cell patterning.

## Results

### *MAKR4* and *MAKR6* are transcriptionally regulated by TIR1/AFB auxin signalling

Our previous transcriptomic analysis identified approximately 125 auxin-induced genes downstream of IAA17/AXR3 and ARF7/19^11,21^ as potential regulators of auxin feedback on PIN polarity. Among these, *MAKR4* and *MAKR6* were also detected in independent datasets^22,23^, but their roles in this context remain unclear.

Using quantitative RT-PCR (qRT-PCR), we confirmed auxin induction of *MAKR4* and *MAKR6*. Specifically, application of 10 µM NAA increased their transcription levels by 52.7- and 4.6-fold, respectively (Fig. 1a, b). Similar induction was observed with 1 µM IAA treatment, but was strongly reduced by 10 µM PEO-IAA (Extended Data Fig. 1a, b), a strong antagonist of TIR1/AFB-mediated auxin perception^24^. These results place *MAKR4* and *MAKR6* downstream of TIR1/AFB signalling. Given the role of WRKY23 in auxin-PIN feedback, we further tested its contribution and found that auxin induction of both genes was markedly reduced in the *wrky23* dominant-negative mutant^11^ (Fig. 1c, d), indicating that WRKY23 regulates their full induction. Therefore, these results position *MAKR4* and *MAKR6* as auxin-inducible genes within the Aux/IAA-ARF pathway and partial targets of WRKY23.

**Fig. 1.**
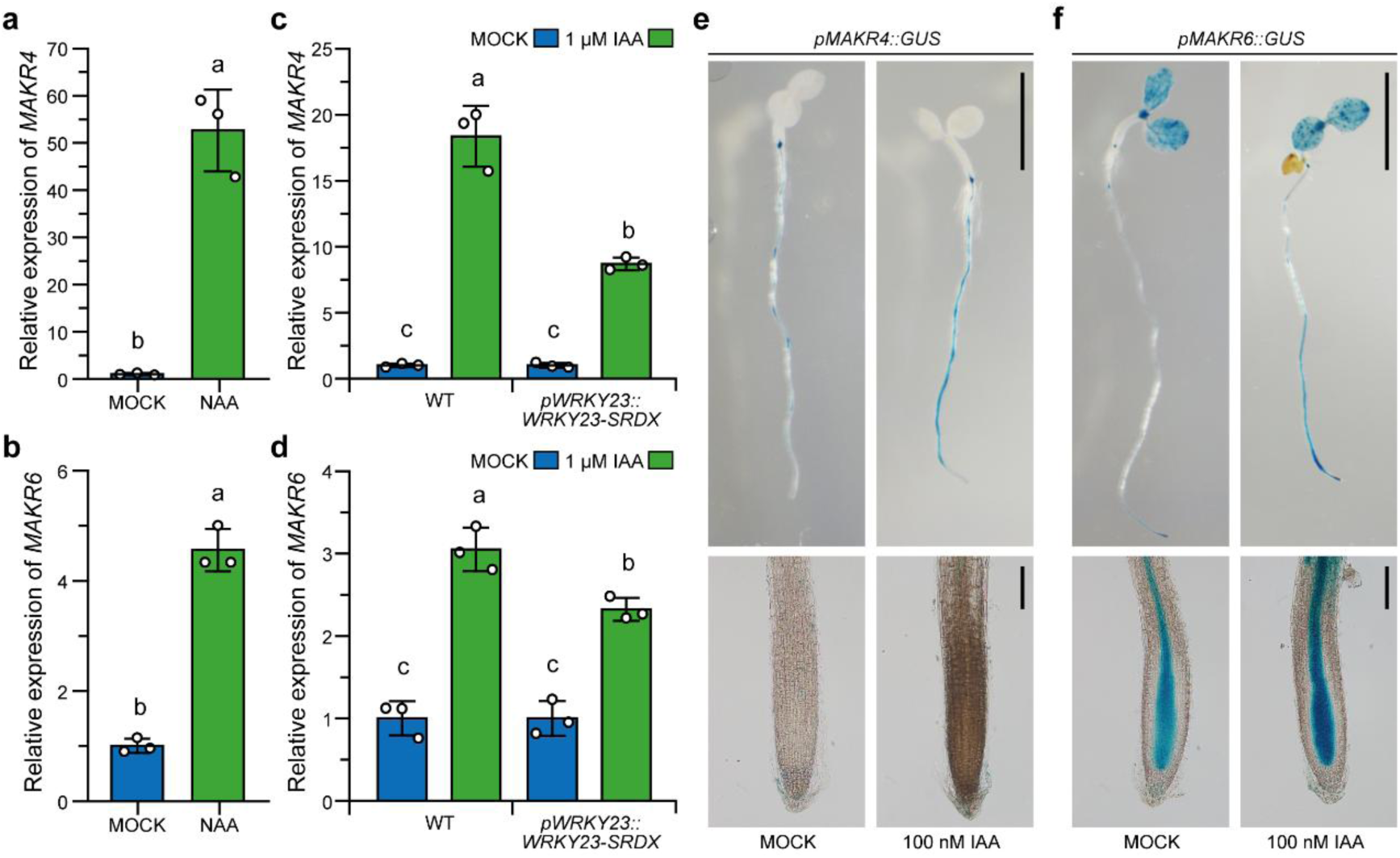
*MAKR4* and *MAKR6* are auxin-inducible genes in the Aux/IAA-ARF-WRKY23 transcriptional axis. **a, b** qRT-PCR analyses of *MAKR4* and *MAKR6* transcription. 5-day-old seedlings were incubated for 4 hours with 10 μM NAA or MOCK before RNA extraction. Relative expression levels were normalized to MOCK controls. **c, d** Auxin induction on the expression of *MAKR4* and *MAKR6* in WT and *wrky*23 mutant. 5-day-old seedlings were incubated with 1 μM IAA or MOCK for 4 hours before RNA extraction. *pWRKY23::WRKY23-SRDX* represents a dominant-negative version of WRKY23 fused to the SRDX transcriptional repression domain. Relative expression levels were normalized to corresponding MOCK controls. **e, f** Expression patterns of *MAKR4* and *MAKR6* in plants. 4-day-old plants of *pMAKR4::GUS* (e) and *pMAKR6::GUS* (f) were used, and the GUS staining was performed after a 4-hour treatment of MOCK (left panels) or 100 nM IAA (right panels). Similar qRT-PCR results were obtained from at least 3 biological replicates. For GUS staining, consistent expression patterns were observed in at least three independent lines, with more than 15 plants analyzed per line. Different letters indicate statistical differences (a, b, Student’s t-test; c, d, two-way ANOVA with Tukey’s HSD, P<0.05). Exact P values for all comparisons are provided in Source Data 1. Scale bars: 5 mm (e, f, upper panels), and 100 μm (e, f, lower panels).

GUS staining of *pMAKR4::GUS* and *pMAKR6::GUS* lines revealed distinct spatial expression patterns. *MAKR4* was mainly expressed in lateral root primordia and mature vascular tissues, while *MAKR6* showed additional expression in vascular tissues from the root tip onward. In leaves, *MAKR6* showed robust expression in cotyledon vasculature and tips, whereas *MAKR4* expression was undetectable. Similarly, *MAKR6*, but not *MAKR4*, showed strong expression in young stems, with apparent induction by wounding (Extended Data Fig. 1c-f). Notably, auxin treatment further enhanced promoter activity and simultaneously expanded expression domains. For instance, *MAKR4* expression became more detectable in mature root regions (Fig. 1e, f, and Extended Data Fig. 1g, h).

Together, these results demonstrate that *MAKR4* and *MAKR6* are auxin-inducible genes in the TIR1/AFB-WRKY23 axis, with distinct expression patterns across tissues.

### MAKR4 and MAKR6 have distinct developmental roles

Compared with WT, *p35S::MAKR4-GFP* and *p35S::MAKR6-GFP* gain-of-function mutants exhibited pronounced defects in primary root growth, lateral root number, and leaf vein patterning (Extended Data Fig. 2a-c), indicating their potential developmental functions. Next, we generated independent CRISPR/Cas9 loss-of-function mutants (*makr4-C1*/*C2* and *makr6-C1*/*C2*), all carrying premature stop codons, and the *makr4-C1;makr6-C1* double mutant by crossing (Extended Data Fig. 2d, e1). Root phenotyping revealed comparable primary root length among WT and all mutants (Extended Data Fig. 2f, g), indicating that neither gene is essential for root elongation. However, lateral root number was significantly reduced in *makr4* but unchanged in *makr6*, as compared with WT (Fig. 2a, b). This indicates that MAKR4 regulates lateral root development, consistent with previous findings^19^. The double mutant phenocopied *makr4* (Fig. 2a, b), suggesting that MAKR4, but not MAKR6, functions primarily in this context.

**Fig. 2.**
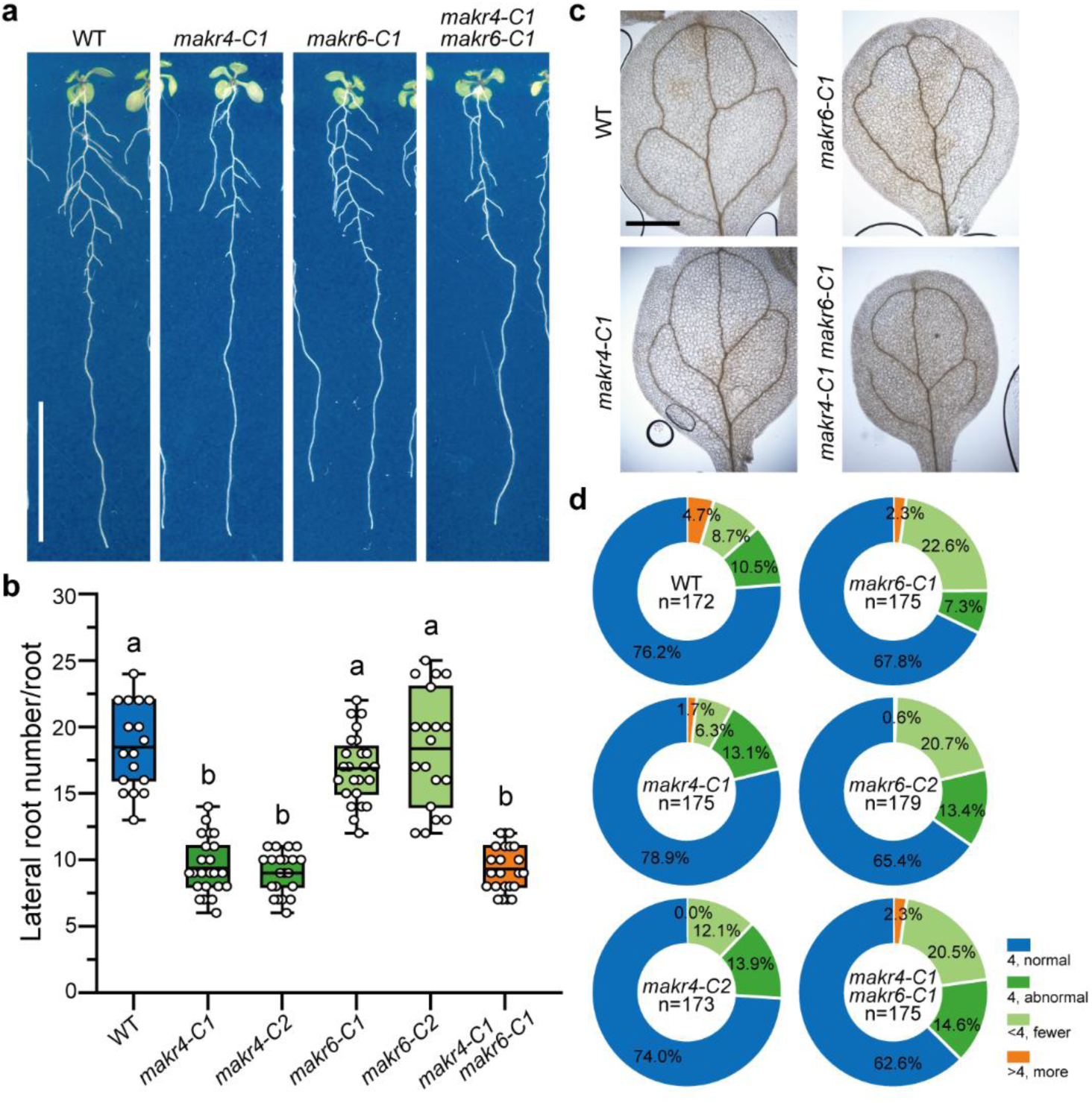
MAKR4 and MAKR6 function distinctly in development. **a, b** Lateral root number in 10-day-old *makr4*, *makr6*, and *makr4;makr6* mutants. Representative images (a) and quantification results of total lateral root number per root (b) are shown. Box plots show the interquartile range (boxes), mean (central lines), and whiskers (min-max values) (n>17 plants per group). Open circles denote individual data points. **c, d** Leaf venation patterns in different genotypes. Representative images (c) and quantification of venation patterns (d) are shown, with sample sizes indicated. 10-day-old plants were used. Different letters indicate statistical differences (b, one-way ANOVA with Tukey’s HSD, P<0.05). Exact P values for all comparisons are provided in Source Data 1. Scale bars: 2 cm (a) and 0.1 cm (c).

Given that *MAKR4* and *MAKR6* act downstream of the SCF^TIR1^-Aux/IAA-ARF and WRKY23 pathway, we next examined their roles in vein vascular patterning, a process closely linked to auxin canalization^5,6,25^. In 10-day-old seedlings, most cotyledons displayed normal four-loop vein patterns in WT (76.2%), *makr4-C1* (78.9%), and *makr4-C2* (74.0%). By contrast, *makr6* mutants showed increased venation defects, with 67.8% of *makr6-C1* and 65.4% of *makr6-C2* cotyledons displaying normal patterns (Fig. 2c, d). These defects were rescued by *pMAKR6::MAKR6-GFP*, confirming that loss of MAKR6 causes the phenotype and that the MAKR6-GFP fusion is functional (Extended Data Fig. 2h). Strikingly, the double mutant phenocopied *makr6* (Fig. 2c, d), indicating that MAKR6, other than MAKR4, predominantly regulates venation patterning.

Together, these results reveal distinct functions: MAKR4 primarily regulates lateral root development, whereas MAKR6 predominantly controls vein patterning. This divergence is consistent with their distinct expression patterns and evolutionary separation into phylogenetic clades (Extended Data Fig. 2i).

### MAKR6 is a novel regulator of vasculature regeneration and formation

To directly test whether MAKR6 acts in auxin canalization, we performed two classical assays: vasculature regeneration after wounding and *de novo* auxin channel formation in response to the exogenous auxin application^1,4,5^.

In the regeneration assay, 90% of WT wounded inflorescence stems formed continuous vascular strands, reconnecting pre-existing vasculature, as revealed by Toluidine Blue O (TBO) staining^12,13^. A similar overall regeneration was observed in *makr4-C1*, although a portion of stems displayed only partial regeneration (60% full, 30% partial) (Fig. 3a, b). However, both *makr6-C1* and *makr4-C1;makr6-C1* mutants rarely formed continuous vascular strands; instead, they exhibited either incomplete vasculature with few lignified cells or no regenerated vessels (Fig. 3a, b). To assess auxin-induced vascular formation, we applied 10 μM IAA below the wound site to induce *de novo* vasculature formation^9,12^. TBO staining showed that most WT and *makr4-C1* stems developed new vascular strands connecting the auxin source to pre-existing vasculature (Fig. 3c, d). However, *makr6-C1* and *makr4-C1;makr6-C1* mutants largely failed to establish such connections, displaying only partial or no vascular development (Fig. 3c, d).

**Fig. 3.**
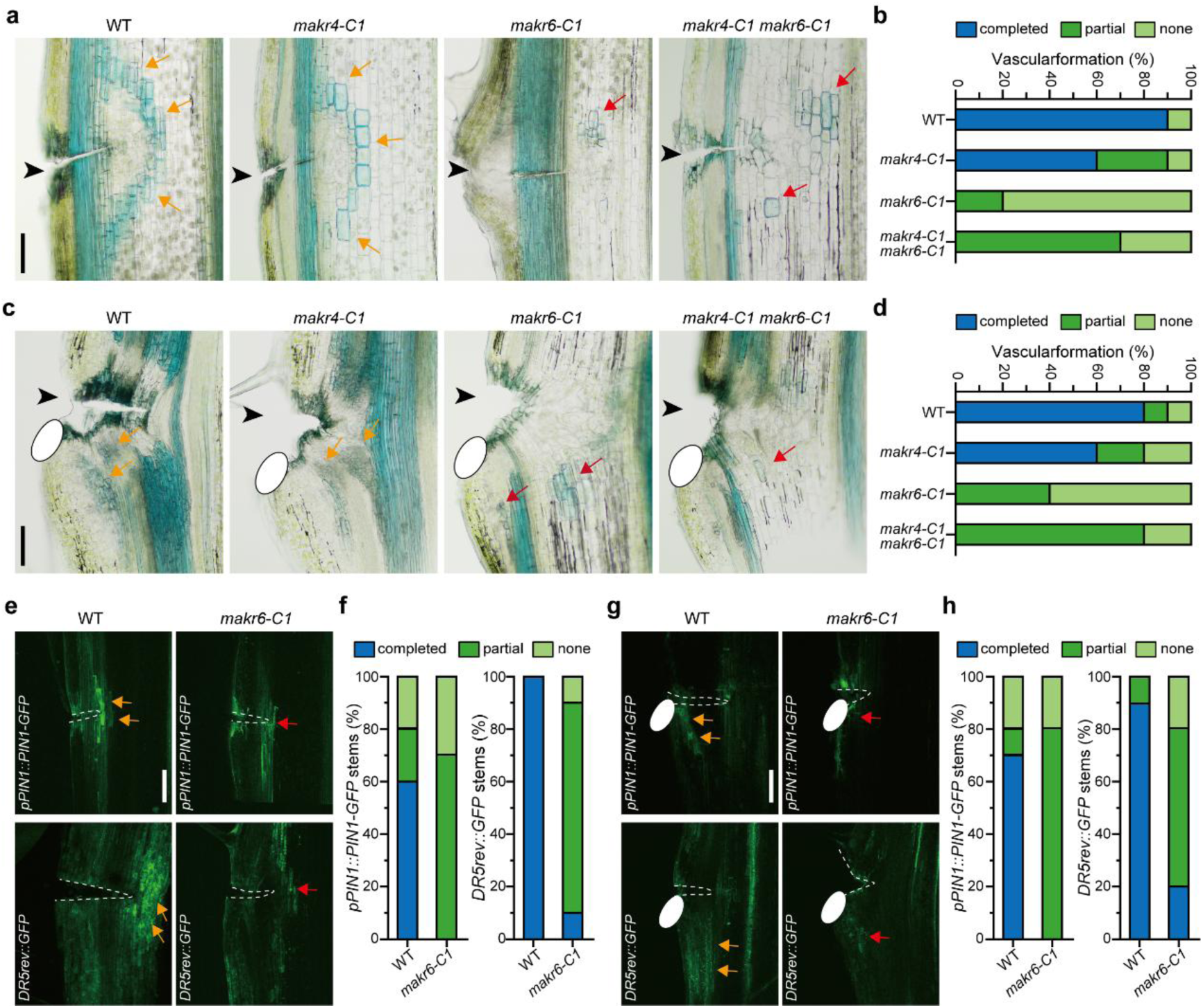
MAKR6 predominantly regulates vascular regeneration and canalization. **a, b** TBO staining of vasculature regeneration after wounding in WT, *makr4-C1*, *makr6-C1*, and *makr4-C1*;*makr6-C1* mutants. Representative images (a) and quantification (b) of vascular regeneration are shown. **c, d** *de novo* vasculature formation induced by exogenous auxin application. Representative images (c) and quantification (d) of vascular regeneration are shown. **e-h** Auxin channel formation visualized by *PIN1::PIN1-GFP* and *DR5rev::GFP*. Panels (e, f) illustrate vascular regeneration, while panels (g, h) show *de novo* auxin channel formation in response to auxin application. Black arrowheads (a, c) mark the wounding sites; orange arrows (a, c, e, and g) indicate newly developed vessels or vasculature; and red arrows (a, c, e, and g) show defective regeneration. 10 μM IAA droplets (white ovals) were applied to the wounded stems (c, g). n=10 plants for each analyzed genotype. Abbreviations: completed, completely formed vasculature; partial, limited, or partially developed vasculature; none, no vessels formed around the wound. Scale bars: 100 μm (a, c, e, and g).

Together, these results establish MAKR6 as an important regulator of both wound-induced regeneration and auxin-driven vascular formation. The similar phenotypes of *makr6-C1* and *makr4-C1;makr6-C1* further support MAKR6 as the primary regulator in these contexts.

### MAKR6 mediates the formation of auxin channels in canalization

Vasculature formation and regeneration are preceded by the establishment of auxin transport channels^5,10^. To examine this process, we introduced the auxin transport marker *pPIN1::PIN1-GFP* and the auxin response marker *DR5rev::GFP*^3,26^ into *makr6-C1*, allowing the monitoring of newly formed auxin channels^9,10^. In vasculature regeneration assays, most WT plants formed complete (60%) or partial (20%) PIN1-GFP channels and continuous *DR5rev::GFP* channels (100%), circumventing wounds (Fig. 3e, f). In contrast, *makr6-C1* mutants rarely formed complete GFP-positive channels in either reporter background (Fig. 3e, f), indicating severe defects in auxin channel initiation and progression. Similar defects were observed in the canalization assay following local auxin application. *makr6-C1* mutants showed strongly impaired auxin channel formation as visualized by *pPIN1::PIN1-GFP* (0%) or *DR5rev::GFP* (20%) markers (Fig. 3g, h). These results demonstrate that MAKR6 mediates self-organized auxin channel formation during the early stages of auxin canalization.

### MAKR6 regulates auxin-dependent PIN polarity and trafficking

The feedback between auxin transport and PIN polarity is central to auxin canalization and can be monitored by auxin-induced PIN1 lateralization and trafficking in roots^5,27^. We therefore tested whether MAKR6 functions in this process. Anti-PIN1 immunostaining showed that in WT roots, treatment with 10 µM NAA induced efficient PIN1 lateralization, with the ratio increasing from 0.79 to 1.13. In contrast, this auxin-induced repolarization was significantly impaired in *makr6-C1* (0.80 to 0.95) (Fig. 4a, b). This indicates that MAKR6 is required for efficient auxin-driven PIN1 repolarization. Next, we applied Brefeldin A (BFA), which blocks PIN1 endomembrane trafficking and induces “BFA bodies”^5,28^. Under MOCK conditions, BFA body formation was comparable between WT and *makr6*. Auxin treatment (10 µM NAA) strongly suppressed BFA body formation in WT roots, including size and density per cell, whereas this inhibition was less pronounced in *makr6* (Extended Data Fig. 3a-c). This suggests that MAKR6 is required for auxin-dependent PIN1 trafficking.

**Fig. 4.**
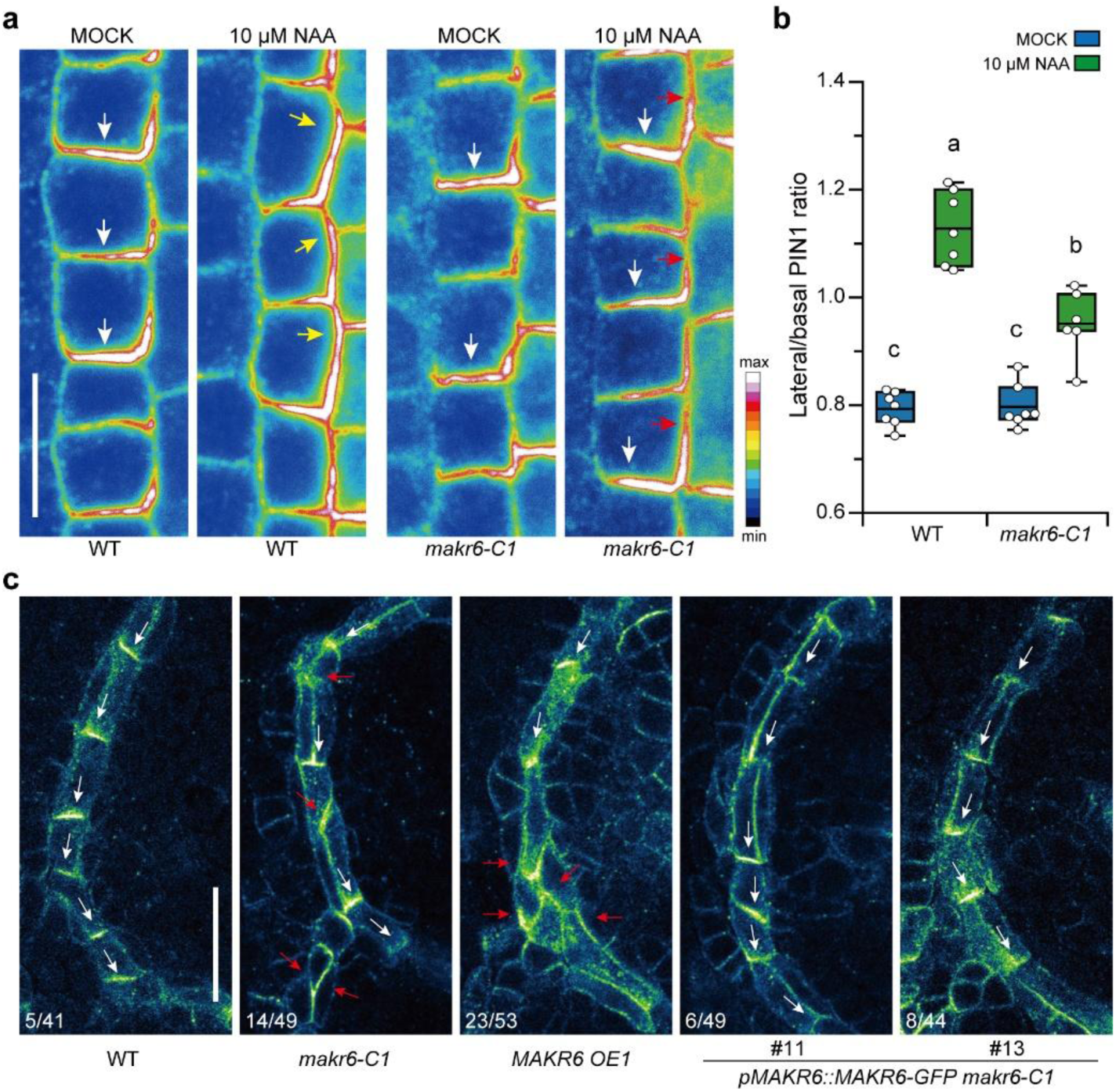
MAKR6 is required for auxin-sensitive PIN1 repolarization and trafficking. **a, b** Auxin-induced PIN1 lateralization in WT and *makr6* roots assessed via anti-PIN1 immunostaining. Representative images of endodermal cells (a) and quantification results (b) are shown. PIN1 lateralization was calculated as the ratio of the inner lateral signal to the rootward basal signal. White and yellow arrows show the predominant PIN1 locations, while red ones indicate abnormal PIN1 accumulation. Data were collected from over 200 cells across at least 7 roots per genotype. **c** Representative images of PIN1 immunolocalization in leaf veins. First leaves of 5-day-old seedlings were analyzed. White arrows indicate typical PIN1 polarity, whereas red arrows indicate altered PIN1 polarity. Numbers indicate the number of leaves displaying PIN1 polarity defects relative to the total number of samples analyzed. Box plot (b) displays the interquartile range (boxes), mean (central lines), and min-max range (whiskers). Open circles denote the averaged result in individual roots. All measurements were performed in a blinded manner, and similar results were obtained from at least two biological replicates. Different letters indicate statistical differences (b, two-way ANOVA with Tukey’s HSD, P<0.05). Exact P values for all comparisons are provided in Source Data 1. Colour scale (a) shows the distribution of the anti-PIN1 signal. Scale bars: 10 μm (a) and 20 μm (c).

Immunolocalization in first leaves showed that WT plants rarely exhibited PIN1 polarity irregularities (5/41 leaves) (Fig. 4c). However, both MAKR6 loss- and gain-of-function lines displayed a significantly higher frequency of defects (14/49 and 23/53 leaves, respectively) (Fig. 4c). This result suggests that MAKR6 is required for robust PIN1 polarity in developing veins.

Notably, these cellular defects were restored in complementation lines (Fig. 4c and Extended Data Fig. 3d, e). Together, these results show that MAKR6 regulates auxin-induced PIN1 repolarization, auxin-sensitive trafficking, and coherent PIN polarity during vein formation, thereby reinforcing the auxin-PIN feedback.

### MAKR6 interacts with TMK1/4 and CAMEL-CANAR receptor-like kinases

Using *pMAKR6::GUS* and *pMAKR6::nlsGFP* reporters, we found that *MAKR6* was predominantly expressed in vascular tissues, with detectable signals in surrounding ground tissues in the root tip. Notably, its expression could be enhanced by auxin while maintaining a similar spatial pattern (Extended Data Fig. 4a-c). Although GFP signal in *pMAKR6::MAKR6-GFP* was below reliable detection, clear PM localization of MAKR6 was observed in *p35S::MAKR6-GFP* and *p35S::TMK1-GFP;p35S::MAKR6-GFP* lines (Extended Data Fig. 4b, d, e).

Considering that BKI1/MAKR proteins frequently act as RLK interactors^16^ and that TMK1/4 and CAMEL/CANAR regulate auxin canalization^12,13^, we asked whether MAKR6 interacts with them. Co-immunoprecipitation (Co-IP) in *Nicotiana* showed that MAKR6 associated with TMK1 and TMK4 (Fig. 5a). Förster resonance energy transfer combined with fluorescence-lifetime imaging microscopy (FRET/FLIM) in Arabidopsis root protoplasts further confirmed these interactions. MAKR6-mNeonGreen (MAKR6-mNG) colocalized with TMK1/4-mCherry (TMK1/4-mCh) at the PM, and co-expression significantly reduced MAKR6-mNG lifetime (Fig. 5b, c). No interaction was observed with the negative control FLS2 (Fig. 5b, c), supporting the interaction specificity. Similarly, direct and specific interactions were detected between MAKR6 and CAMEL/CANAR by both Co-IP and FRET/FLIM (Fig. 5d-f). Auxin treatment did not noticeably alter these interactions, as comparable reductions in MAKR6-mNG lifetime were observed under NAA and MOCK conditions (Extended Data Fig. 5a).

**Fig. 5.**
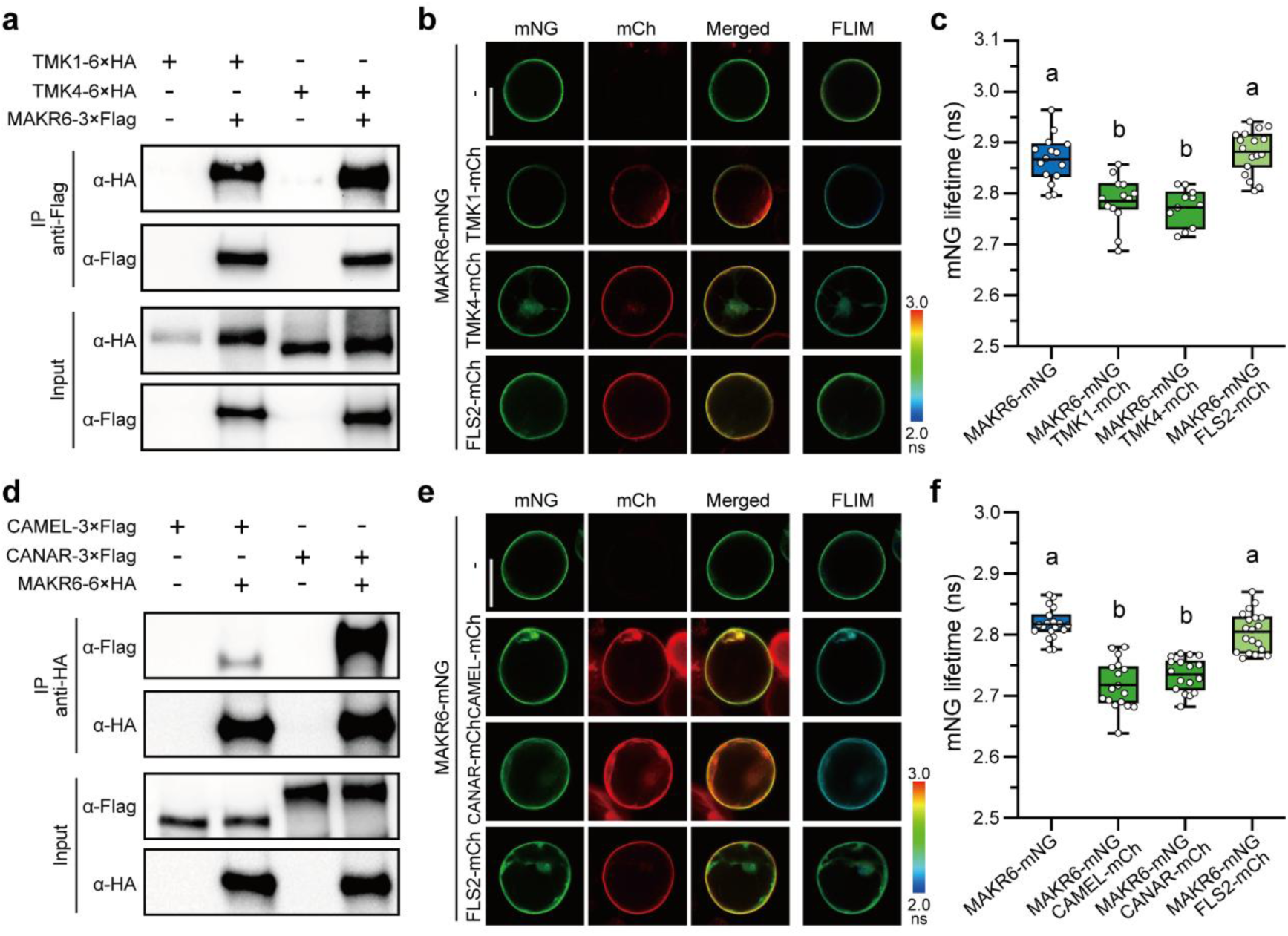
MAKR6 interacts with TMK1/4 and CAMEL/CANAR. **a** Co-IP assay from Nicotiana leaves transiently expressing TMK1/4-6×HA alone or with MAKR6-3×Flag. Immunoprecipitation was performed using anti-Flag beads, and the samples were analyzed by Western blot. **b, c** FRET/FLIM in Arabidopsis root protoplasts transformed with *p35S::MAKR6-mNG* and *p35S::TMK1/4-mCh* constructs. Representative images (b) and the mNG lifetime quantification (c), with data collected from more than 11 cells per group. *p35S::FLS2-mCh* was used as a control. **d-f** Interaction between MAKR6 and CAMEL/CANAR determined by the Co-IP from Nicotiana leaves (d) and FRET/FLIM inArabidopsis (e, f). Co-IP was performed using anti-HA beads, and FRET/FLIM analysis was performed on more than 17 cells per group. Box plots (c, f) show the interquartile range (boxes), mean (central lines), and min-max range (whiskers) of the fluorescence lifetime measured in individual cells (open circles). Each experiment was independently replicated at least three times, yielding similar results; one representative replicate is shown. Abbreviations: mCh, mCherry; mNG, mNeonGreen. Different letters indicate statistical differences (c, f, one-way ANOVA with Tukey’s HSD, P<0.05). Exact P values for all comparisons are provided in Source Data 1. The colour bars (b, e) indicate the distribution of mNG fluorescence lifetimes. Scale bars: 30 μm (b, e).

Collectively, these results identify MAKR6 as a PM-associated interactor of TMK1/4 and CAMEL/CANAR, supporting a role for MAKR6 in integrating RLK signalling to auxin-dependent canalization.

### MAKR6 is phosphorylated by associated RLKs acting on PIN1 dynamics and leaf venation

As both TMKs and CAMEL possess kinase activity^12,13,29,30^, we examined whether they could phosphorylate MAKR6. Phos-tag analyses revealed a pronounced mobility shift of MAKR6 when co-expressed with TMK1/4 or CAMEL in protoplasts (Extended Data Fig. 5b, c), indicating that MAKR6 is phosphorylated by these kinases. Together with the previously described MAKR2-TMK1 and BKI1-BRI1 pairs^16,18,31^, our observations support a conserved phosphorylation regulatory mechanism in MAKR-RLK modules.

To define functional connections, we generated higher-order mutants by crossing *makr6* with *tmk4*, *camel*, and *canar irk*^32^ mutants, respectively. Phenotypic analyses showed that auxin-induced PIN1 repolarization and trafficking defects in roots were comparable between higher-order mutants and *makr6-C1* (Extended Data Fig. 5d, e). Consistently, combined mutants did not display enhanced leaf venation defects relative to *makr6 or corresponding rlk mutants* (Extended Data Fig. 5f). These genetic interactions indicate that MAKR6, TMKs, and CAMEL/CANAR function within a shared signalling pathway.

Together, these results indicate that MAKR6 integrates inputs from CAMEL- and TMK-mediated pathways, likely through phosphorylation-dependent regulation. The output converges on auxin-PIN1 feedback, thereby contributing to auxin canalization.

### TMK1/4 associate with the CAMEL-CANAR complex

The genetic convergence of TMK1/4 and CAMEL-CANAR prompted us to test whether these RLKs interact. Co-IP assays in *Nicotiana* demonstrated that both CAMEL and CANAR were pulled down with either TMK1-3×Flag or TMK4-3×Flag (Fig. 6a, b), suggesting physical associations. FRET/FLIM analyses in protoplasts further validated these interactions, as co-expression of TMK1/4-mCh significantly reduced GFP lifetime of CAMEL-GFP or CANAR-GFP (Fig. 6c-f). Notably, auxin showed no detectable effect on the TMK1-CAMEL interaction (Extended Data Fig. 6a). Given that TMK1/4 and CAMEL/CANAR also interact with MAKR6, we next asked whether MAKR6 associates with or mediates this complex. FRET/FLIM between TMK1-mNG and CAMEL/CANAR-mCh was performed in the presence of MAKR6-mTurquoise2 (MAKR6-mTQ2) or FLS2-mTQ2. According to FRET/FLIM principles, incorporation of an additional component into a stable complex typically alters the donor-acceptor spatial geometry and FRET efficiency^33^. However, comparable reductions in FRET efficiency were observed in both conditions (Fig. 6g; Extended Data Fig. 6b), suggesting that MAKR6 does not measurably modulate or associate with the TMK-CAMEL/CANAR complex under the conditions tested.

**Fig. 6.**
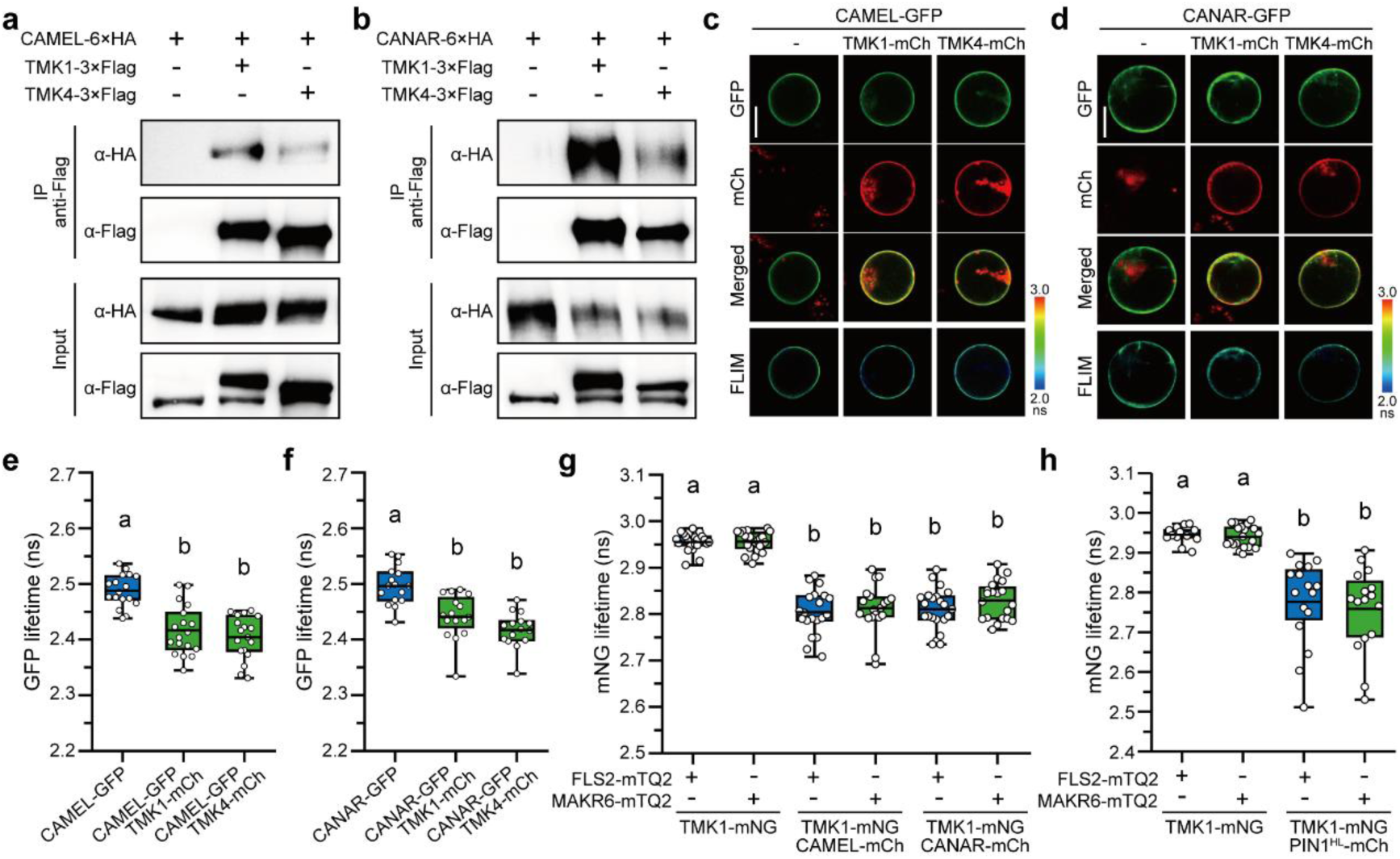
The association between TMK1/4 and the CAMEL-CANAR complex. **a, b** CAMEL-CANAR interaction with TMK1/4, as revealed by Nicotiana Co-IP. Anti-Flag Co-IPs were performed with leaves transiently transformed with *p35S::TMK1/4-3×Flag*, *p35S::CAMEL-6×HA*, and *p35S::CANAR-6×HA* in combinations. The input samples show unspecific anti-Flag bands. **c-f** FRET/FLIM assays showing the interaction between the CAMEL-CANAR complex and TMK1/4. Either *p35S::CAMEL-GFP* or *p35S::CANAR-GFP* was transformed alone or with *p35S::TMK1/4-mCh* in combinations. At least 16 cells were analyzed per group. **g, h** MAKR6 interactions with TMK1 and CAMEL/CANAR and PIN1. FRET/FLIM experiments were done in protoplasts expressing TMK1-mNG alone or with CAMEL/CANAR/PIN1^HL^-mCh in the presence of MAKR6-mTQ2. Lifetime was measured from at least 20 cells (g) and 13 cells (h). Box plots (e-h) show the fluorescence lifetime measured in individual cells (open circles), with boxes indicating the interquartile range, central lines the mean, and whiskers the min-max range. Each experiment was independently replicated at least twice, yielding consistent results; one representative replicate is shown. Abbreviations: mCh, mCherry; mNG, mNeonGreen; mTQ2, mTurquoise2. Different letters indicate statistical differences (e, f, one-way ANOVA; g, h, two-way ANOVA with Tukey’s HSD, P<0.05). Exact P values for all comparisons are provided in Source Data 1. The colour bars (c, d) show the GFP lifetime in the FRET/FLIM. Scale bars, 30 μm (c, d).

Our results identify a TMK-CAMEL/CANAR signalling complex at the cell surface that may integrate auxin and peptide-derived cues during canalization. In contrast, MAKR6 appears to function independently, likely acting dynamically or downstream of these complexes.

### MAKR6 interacts with the PIN1 auxin transporter

Given that MAKR2 controls PIN2 polarity in gravitropism^18^ and MAKR6 regulates PIN1 dynamics in canalization, we asked whether MAKR proteins directly associate with PIN transporters. FRET/FLIM in Arabidopsis protoplasts showed that co-expression of MAKR6-GFP with mCherry-tagged PIN1 hydrophilic loop (PIN1^HL^) significantly reduced GFP lifetime (Extended Data Fig. 7a), indicating a direct MAKR6-PIN1 interaction. Auxin treatment did not noticeably affect this interaction (Extended Data Fig. 5a). The MAKR6-PIN1 interaction was further confirmed using full-length PIN1 in independent FRET/FLIM and Co-IP assays. Neither FLS2 nor MAKR2 exhibited detectable interaction with PIN1 or MAKR6, supporting interaction specificity (Extended Data Fig. 7a-c). Moreover, Arabidopsis Co-IP confirmed these associations *in planta*: HA- or Flag-tagged MAKR6 co-precipitated with endogenous TMK1 and PIN1 (Extended Data Fig. 7d, e).

We next extended this analysis to MAKR4, given its role in lateral root initiation, in which TMK1/4 and PIN1 also function^19,27,34^. FRET/FLIM detected no MAKR4-TMK1/4 interaction but a clear MAKR4-PIN1^HL^ interaction, which was further confirmed using full-length PIN1 (Extended Data Fig. 7f, g). These findings define a distinct MAKR4-PIN1 module, likely operating with other RLK partners besides TMK1/4.

Given that TMKs and CAMEL-CANAR also associate with PIN1^12,30^, we tested whether MAKR6 modulates these interactions, using TMK1-PIN1 as a representative. FRET/FLIM showed identical energy transfer between TMK1-mNG and PIN1^HL^-mCh with MAKR6-mTQ2 or FLS2-mTQ2 (Fig. 6h; Extended Data Fig. 7h). This suggests that MAKR6 does not modulate or associate with the TMK1-PIN1 complex.

Together, these results demonstrate that MAKR4 and MAKR6 associate with PIN1, supporting a role in regulating PIN1 polarity. This may represent a conserved MAKR-PIN regulatory module, potentially extending to MAKR2-PIN2.

## Discussion

This work identifies MAKR6 as a multifaceted regulator of auxin canalization, coupling auxin signalling with PIN polarization and integrating multiple RLK complexes. A model emerging from our findings is that MAKR6 coordinates distinct RLKs to fine-tune auxin canalization (Extended Data Fig. 8). MAKR6 interacts with CAMEL/CANAR, TMK1/4, and PIN1, likely within distinct dynamic complexes. These pathways are integrated into the auxin-PIN1 feedback loop, supporting robust vascular patterning and regeneration. TMK- and CAMEL-associated signalling regulate MAKR6 through phosphorylation, potentially downstream of ligand perception.

Both *MAKR4* and *MAKR6* transcripts are auxin-regulated via the SCF^TIR1^-Aux/IAA-ARF-WRKY23 axis. Despite this shared regulation, they diverge in expression and function. MAKR6 predominantly contributes to classical auxin canalization-dependent processes, whereas MAKR4 primarily regulates lateral root initiation, which also involves dynamic PIN re-arrangements^19,27^. Thus, our work showcases the divergent evolution of BKI1/MAKR proteins through gradual differentiation of function, expression, and protein interaction.

Auxin canalization features a positive feedback loop between auxin signalling and PIN polarity^1,5,6^. Our work defines MAKR6 as a molecular bridge in this loop, filling another mechanistic gap in the canalization model. *MAKR6* is auxin-regulated and contributes to multiple canalization-mediated processes. The direct MAKR6-PIN1 interaction provides a mechanistic basis for its regulation of PIN1 polarity and trafficking, and similar mechanisms may apply to other MAKRs (e.g., MAKR2-PIN2 and MAKR4-PIN1).

Receptor-like kinases function as signal receptors, scaffolds, or nanocluster components at the cell surface^35–37^. We previously identified TMKs, CAMEL, and CANAR as major canalization regulators^12,13^ and now show that all interact with MAKR6. These PM-localized RLKs likely bind their respective cues to mediate intercellular communication within and around the auxin canal. The CAMEL/CANAR-TMK1/4 complex indicates signal convergence, supported by genetic interactions among MAKR6, CAMEL/CANAR, and TMK1/4. This fine-tuned regulation may help ensure precise patterning across timescales and cell contexts. How the MAKR4-PIN1 module and its RLK partners function in lateral root development, another process requiring auxin and PIN repolarization, remains to be determined.

RLKs dynamically interact with partners before, during, and after ligand binding, and BKI1/MAKRs are such regulators^16,35,38^. For instance, BKI1 negatively regulates BR signalling by preventing BRI1-BAK1 association^16,37,39^. MAKR2 negatively regulates TMK1 in root gravitropism^18^, whereas MAKR5 amplifies CLE45-BAM3 signalling in protophloem differentiation^17,20^. Our findings show that MAKR6 coordinates CAMEL-CANAR and TMK1/4 in canalization. However, whether MAKR6 acts positively or negatively remains unclear, as both loss- and gain-of-function mutants exhibit severe canalization defects. MAKR6 is phosphorylated by CAMEL and TMKs, but auxin or other cues may also regulate its localization and interactions in a feedback manner, a direction for future work.

Auxin canalization involves RLKs that may work synergistically, antagonistically, and sequentially^5^. Although CAMEL/CANAR interacts with TMK1/4, MAKR6 does not directly influence these interactions, suggesting it acts independently within each pathway rather than mediating cross-complex assembly. Instead, MAKR6 may modulate each RLK interaction with its co-receptor during ligand perception (Extended Data Fig. 8). The possibility that a single MAKR6 protein acts on multiple RLKs (at least CAMEL-CANAR and TMK1/4) could explain the observed phenotypic complexity. Future studies on specific ligands, co-receptors, and downstream cascades will help elucidate MAKR6’s multifaceted function.

In summary, auxin canalization is a fundamental mechanism driving plant patterning and organogenesis. Our study identifies MAKR6 as an important regulator that integrates multiple intercellular signalling networks and mediates auxin-PIN1 feedback. These discoveries expand our understanding of the molecular components and intricate RLK interplay that orchestrates complex developmental processes.

## Methods

### Plant materials and growth conditions

All experiments were conducted using *Arabidopsis thaliana* Columbia-0 (Col-0) as the wild-type (WT) and transformation background, and *Nicotiana benthamiana* plants for transient expression assays. Seeds were sterilized overnight using chlorine gas (39 mL bleach, 4.5 mL HCl, 61 mL H_2_O) and stratified at 4 °C in the dark for 24 hours before germination. Seedlings were grown on solidified 1/2 Murashige and Skoog (MS) medium under long-day conditions (16-hour light/8-hour dark cycles, 22 ± 2°C, 100 μmol m⁻² s⁻¹). Seedlings were cultivated vertically for 4-10 days, depending on the experiment. After germination, seedlings were transferred to soil and grown under the same photoperiod and temperature conditions.

Transgenic lines and mutants generated in this study included *pMAKR4::GUS*, *pMAKR6::GUS*, *pMAKR6::nlsGFP*, *p35S::MAKR4-GFP*, *p35S::MAKR6-GFP*, *p35S::MAKR6-6 × HA*, *pMAKR6::MAKR6-GFP*, *p35S::MAKR6-3 × Flag*, *p35S::TMK1-GFP;p35S::MAKR6-mCherry*, *makr4-C1*, *makr4-C2*, *makr6-C1*, *makr6-C2*, and *pMAKR6::MAKR6-GFP makr6-C1*. Previously described reporter lines and mutants included *pWRKY23::WRKY23-SRDX*, *DR5rev::GFP*, *PIN1::PIN1-GFP*, *camel-1*, *tmk4*, and *canar irk*^3,11,12,26,32,40^. Higher-order mutants and reporter combinations were generated by genetic crossing. All plant materials were verified by PCR genotyping, sequencing, and/or antibiotic selection before use.

### Plasmid construction and plant transformation

Gateway cloning (Thermo Fisher), including pENTR/D-TOPO entry cloning, BP reactions, and LR recombination, was used as the main cloning strategy throughout this study according to the manufacturer’s instructions. Donor and destination vectors, together with the resulting entry and expression constructs, are listed in Supplementary Table 1. Additional destination vectors were obtained from previous studies, including pB7FLAGWG2, pB7GUSWG0, and pB7MWG2^41,42^. Previously generated constructs included CAMEL, CANAR, and MAKR2 entry clones in pDONR221, as well as *p35S::PIN1-GFP*, *p35S::mCh-PIN1^HL^*, and *p35S::PIN1^HL^-mCh*^12,18^.

For FRET/FLIM assays of MAKR6 interactions with CAMEL/CANAR/TMK1/4, the following constructs were used: *gMAKR6*/pCoE2GWmNG7, *gTMK1*/pCoE2GWmCh7, *gTMK4*/pCoE2GWmCh7, *CAMEL*/pCoE2GWmCh7, *CANAR*/pCoE2GWmCh7, and *FLS2*/pCoE2GWmCh7.

For FRET/FLIM analyses examining TMK1/4 interactions with CAMEL/CANAR, constructs *CAMEL*/pB7FWG2, *CANAR*/pB7FWG2, *gTMK1*/pB7MWG2, and *gTMK4*/pB7MWG2 were used in the indicated combinations. For FRET/FLIM assays testing interactions between MAKR proteins and PIN1, constructs included *MAKR4*/pB7FWG2, *gMAKR6*/pB7FWG2, *PIN1^HL^*/p2GWmCh7, *PIN1^HL^*/p2mChGW7, *PIN1-GFP*/pK7WG2, *MAKR2*/pB7MWG2, *MAKR4*/pB7MWG2, *gMAKR6*/pB7MWG2, *gTMK1*/pB7MWG2, *gTMK4*/pB7MWG2, and *FLS2*/pB7MWG2.

To assess the effect of MAKR6 on TMK1 interactions in FRET/FLIM assays, *CAMEL*/pCoE2GWmCh7, *CANAR*/pCoE2GWmCh7, *PIN1^HL^*/p2GWmCh7, *gMAKR6*/pCoE2GWmTQ27-*gTMK1*/p2GWmNG7, and *FLS2*/pCoE2GWmTQ27- *gTMK1*/p2GWmNG7 were used.

For auxin-treatment FRET/FLIM assays, constructs included *gMAKR6*/pCoE2GWmNG7, *CAMEL*/pCoE2GWmCh7, *gTMK1*/pCoE2GWmCh7, *PIN1^HL^*/p2GWmCh7, *gTMK1*/pB7FWG2, and *CAMEL*/pB7MWG2.

For Phos-tag assays in protoplasts, *gMAKR6*/pCoE2GWmTQ27, *gMAKR6*/pCoE2GWmTQ27-*gTMK1*/p2GWmNG7, *gMAKR6*/pCoE2GWmTQ27-g*TMK4*/p2GWmNG7, and *gMAKR6*/pCoE2GWmTQ27-*CAMEL*/p2GWmNG7 were used.

### GUS staining and qRT-PCR

Seedlings or young shoots of GUS reporter lines, with or without 4-hour auxin pre-treatments, were harvested and incubated in ice-cold 90% acetone for 30 min. Samples were subsequently washed three times with GUS staining buffer (16 mM NaH₂PO₄, 34 mM Na₂HPO₄, 0.5 mM K₄[Fe(CN)₆], 0.5 mM K₃[Fe(CN)₆]) and incubated at 37 °C in 2 mM X-Gluc solution. Staining duration was adjusted according to promoter activity. Reactions were terminated by replacing the staining solution with 70% ethanol for tissue clearing. Finally, the GUS-stained seedlings were mounted in water and imaged using an Olympus BX53 differential interference contrast microscope and an Olympus SZX16 stereo microscope.

qRT-PCR analyses were performed as described previously^11,12^. Briefly, 5-day-old whole seedlings were used for RNA extraction after 4-hour treatments in AM medium, as specified in each experiment. Total RNA was extracted and used for cDNA synthesis, followed by quantitative PCR analysis. Relative expression levels were calculated using the 2^-ΔΔCt method, normalized to *PP2A* expression, and further normalized to MOCK controls.

### Root phenotypic analyses

For root growth measurements, seedlings were grown vertically and scanned 6 days post-germination. Root length was measured using the SNT plugin^43^ in ImageJ (http://rsb.info.nih.gov/ij). Similarly, 4-day-old vertically grown seedlings were transferred to fresh plates, and six days post-transfer, the total number of lateral roots was visually counted from the scanned images.

### PIN1 immunostaining

Auxin-induced PIN1 lateralization and trafficking in roots were analyzed by whole-mount immunolocalization as described previously^27,44^. Briefly, 4-day-old seedlings were incubated in liquid 1/2 MS medium containing 10 μM NAA or MOCK (DMSO) for 4 hours to assess PIN1 lateralization, or in medium containing 50 μM BFA together with 10 μM NAA or MOCK for 2 hours to examine PIN1 trafficking. After incubation, the samples were fixed in 4% paraformaldehyde (PFA) and immunostained with an anti-PIN1 antibody, followed by a Cy3-conjugated goat anti-rabbit secondary antibody (Sigma-Aldrich).

PIN1 immunolocalization in first leaves was performed as described previously^12^. 5-day-old seedlings were fixed in 4% PFA for 60 min and washed twice in PBS for 10 min each. Samples were sequentially cleared in methanol (3 × 10 min, 37 °C), ethanol:p-xylene (1:1; 3 × 10 min, 37 °C), p-xylene (3 × 10 min, 37 °C), and ethanol:p-xylene (1:1; 3 × 10 min). Seedlings were then rehydrated through a graded ethanol series (100%, 90%, 75%, 50%, and 25%) for 5 min each, followed by H_2_O washes before immunolocalization.

Imaging was performed using Zeiss LSM800, LSM880, and LSM900 confocal microscopes with Cy3 detection settings, and image analyses were conducted using ImageJ. Quantification of PIN1 lateralization and BFA body formation was performed in endodermal and pericycle cells, respectively, where signals were consistently distinguishable and quantifiable. 16-colour lookup tables were applied for PIN1 lateralization analyses, Magenta LUTs for BFA-treated samples, and Green Fire Blue LUTs for PIN1 polarity analyses in leaves.

### Cotyledon vasculature analysis

Leaf vasculature analysis was performed using cotyledons from 10-day-old seedlings of the indicated genotypes. After root removal, shoot tissues were fixed in ethanol:acetic acid solution (3:1) for 1 hour. Following three rounds of fixation, samples were cleared in 70% ethanol before imaging. Cotyledons were subsequently dissected, mounted in water, and imaged using an Olympus BX53 differential interference contrast microscope.

Vein patterns were classified based on loop connectivity and loop number. Cotyledons containing four fully connected loops, including those with basal loops open at the bottom, were classified as “4 loops, normal”. Cotyledons with disruptions at other positions within the four-loop pattern were categorized as “4 loops, abnormal”. Those with fewer than four loops (<4) or more than four loops (>4), regardless of additional defects, were categorized as “fewer loops” or “more loops” respectively.

### Phylogenetic analysis

Full-length protein sequences of BKI1 and MAKR1-6 were obtained from The Arabidopsis Information Resource (TAIR) (www.arabidopsis.org). A neighbour-joining phylogenetic tree was constructed using the Bootstrap method and Poisson substitution model in MEGA 12.0 (www.megasoftware.net).

### Vasculature regeneration after wounding and *de novo* auxin canalization

Vasculature regeneration after wounding was performed as described previously^12,44^. Briefly, inflorescence stems of young Arabidopsis plants (∼10 cm tall) were transversely incised above the rosette leaves to disrupt vascular continuity and polar auxin transport. Regenerated vasculature was analyzed 6 days after wounding (DAW) using 0.05% Toluidine Blue O (TBO) staining. For *de novo* auxin canalization assays, lanolin paste containing 10 μM IAA was applied below the incision site and replaced every 2 days. Regenerated vasculature was analyzed 6 days after auxin application (DAA) using TBO staining. Sections were mounted in 50% aqueous glycerol and imaged using an Olympus BX4 bright-field microscope.

To monitor auxin channel formation, the reporter lines *pPIN1::PIN1-GFP* and *DR5rev::GFP* were introduced into the indicated genetic backgrounds. GFP fluorescence was imaged 4 days after wounding using an Olympus FLUOVIEW FV1000 confocal microscope. All phenotypic evaluations were performed blindly using coded samples to minimize experimental bias.

### Co-immunoprecipitation assays in plants

Co-IP assays in *Nicotiana* leaves were performed following Agrobacterium-mediated transient expression as described previously^41,45^. Constructs encoding tagged proteins were transformed into *Agrobacterium tumefaciens GV3101* and infiltrated into young leaves individually or in the indicated combinations. Following overnight incubation in the dark and subsequent growth for 24 hours in greenhouse conditions, leaves were harvested and homogenized on ice in lysis buffer (50 mM Tris-HCl, pH 8.0, 150 mM NaCl, 10% glycerol, 1 mM PMSF, 5 mM DTT, 0.5% NP-40, and 1×cOmplete protease inhibitor). Lysates were clarified by two rounds of centrifugation and incubated with Pierce™ Anti-DYKDDDDK Magnetic Agarose or Pierce™ Anti-HA Magnetic Beads (Thermo Fisher) for 2 hours at 4 °C with gentle rotation. Immunocomplexes were washed extensively, separated by SDS-PAGE, and detected using anti-HA-Peroxidase High Affinity (Sigma-Aldrich), anti-Flag-HRP (Sigma-Aldrich), or anti-GFP-HRP (Miltenyi Biotec) antibodies.

For Co-IP assays in Arabidopsis, 8-day-old seedlings were ground in liquid nitrogen and homogenized in extraction buffer (25 mM Tris-HCl, pH 7.5, 150 mM NaCl, 0.1% Triton X-100, 0.5 mM DTT, 1×cOmplete protease inhibitor) at 2 μL/mg tissue. Extracts were incubated on ice for 20 min with intermittent vortexing, then immunoprecipitated and immunoblotted as described above. Endogenous PIN1 and TMK1 were detected using previously described antibodies^27,46^.

### FRET/FLIM in Arabidopsis root suspension culture protoplasts

FRET/FLIM experiments were performed in Arabidopsis root protoplasts using PEG-mediated transfection, following established protocols^42,47^. Protoplasts were isolated from 3-day-old suspension cultures via 4 hours of enzymatic digestion (1% Cellulase RS, 0.2% Driselase, and 0.05% Pectinase) in the B5-0.34M glucose-mannitol (GM) solution (2.2 g MS with vitamins, 16.77 g glucose, 15.25 g mannitol, H_2_O to 500 mL, pH 5.5). Following centrifugation and washing, cells were floated in B5-0.28 M sucrose solution (0.44% MS with vitamins, 0.28 M sucrose, pH 5.5). The final cell density was adjusted to 4×10⁶ cells/mL in GM solution.

For transfection, 50 μL protoplast suspension was mixed with 10 μg plasmid DNA and 200 μL PEG buffer (0.1 M Ca(NO_3_)_2_, 0.45 M D-mannitol, and 25% PEG 6000, pH 9.0). After 30 min of incubation on ice in the dark, protoplasts were washed twice with 0.275 M Ca(NO₃)₂ and incubated overnight in GM solution at room temperature. For auxin-treatment experiments, 300 nM NAA in GM solution was added to protoplasts at a 1:2 dilution and incubated for 2 hours before imaging.

Transfected protoplasts were transferred to Petri dishes and imaged using a Leica SP8 microscope equipped with an FLIM module. Fluorophores, including mTurquoise2, mNeonGreen, GFP, and mCherry, were detected using the corresponding excitation and emission settings. Fluorescence lifetime measurements were analyzed with LAS X software (FLIM).

### Phos-tag assays in Arabidopsis protoplasts

Phos-tag assays were performed using Arabidopsis protoplasts prepared and transfected as described above. Protoplasts expressing the indicated protein combinations were collected by centrifugation. Pellets were resuspended in 100 μL extraction buffer (25 mM Tris-HCl, pH 7.6, 150 mM NaCl, 0.1% Triton X-100, 1×cOmplete protease inhibitor cocktail, and 1×PhosSTOP phosphatase inhibitor). Samples were rotated at 4 °C for 30 min, centrifuged to collect the supernatant, mixed with sample buffer (Wako), and heated at 98 °C for 2 min. For λ protein phosphatase (λ-PP) treatment, 15 μL of protein lysate was incubated with 5 μL of λ-PP reaction components (NEB) at 30 °C for 1 hour, followed by termination with sample buffer.

Phos-tag SDS-PAGE gels were prepared essentially as described previously^48^. Resolving gels (4.5%) containing 5 mM Phos-tag acrylamide (Wako) and 20 mM ZnSO₄ were cast, followed by stacking gels after polymerization. Protein samples were separated by electrophoresis, and gels were incubated in washing buffer (25 mM Tris- HCl, 200 mM glycine, 5% methanol, 10 mM EDTA) for 30 min before wet transfer. Proteins were detected by immunoblotting using anti-mNeonGreen (Agrisera), anti-rabbit secondary antibody-HRP (Thermo Fisher), or anti-GFP-HRP (Miltenyi Biotec).

### Statistical analyses

Statistical analyses were performed using Student’s t-test or ANOVA in GraphPad Prism (www.graphpad.com). Graphs and data plots were generated with OriginPro 2022 (www.originlab.com), and figures were assembled using Adobe Photoshop 2025 and Adobe Illustrator 2025 (www.adobe.com).

## Data availability

All data supporting the findings of this study are available from the corresponding author upon reasonable request. Key resource data are provided with this paper and supplementary materials. Gene accession numbers used in this study are as follows: BKI1 (AT5G42750), MAKR1 (AT5G26230), MAKR2 (AT1G64080), MAKR3 (AT2G37380), MAKR4 (AT2G39370), MAKR5 (AT5G52870), MAKR6 (AT5G52900), CAMEL (AT1G05700), CANAR (AT5G01890), IRK (AT3G56370), TMK1 (AT1G66150), TMK4 (AT3G23750), FLS2 (AT5G46330), PIN1 (AT1G73590), and PIN2 (AT5G57090).

## Acknowledgments

We would like to thank Dr. Yvon Jaillais (ENS, Lyon) for sharing MAKR2 materials. This research was supported by the Scientific Service Units (SSU) of ISTA through resources provided by the Imaging & Optics Facility (IOF) and the Lab Support Facility (LSF). The research in the Friml group leading to these results was funded by the European Research Council (ERC): 101142681 CYNIPS; and the Austrian Science Fund (FWF): I 6123-B and P 37051-B. Ewa Mazur was supported by the National Science Centre (NCN), Poland, under the OPUS call in the WEAVE programme: 2021/43/I/NZ1/01835.

## Author contributions

Conceptualization, Z. G., G. M., and J. F.; Methodology, Z. G., and J. F.; Investigation, Z. G., L. K., E. M., G. M., S. I., D. V., N. K., and L. F.; Writing-Original Draft, Z. G., and J. F.; Writing-Review & Editing, Z. G., E. M., and J. F.; Funding Acquisition and Resources, E. M., and J. F.; Supervision, Z. G., and J. F.

## Ethics declarations

Competing interests: The authors declare no competing interests.

**Correspondence** and requests for materials should be addressed to Jiří Friml.

## Extended data for

**Extended Data Fig. 1.**
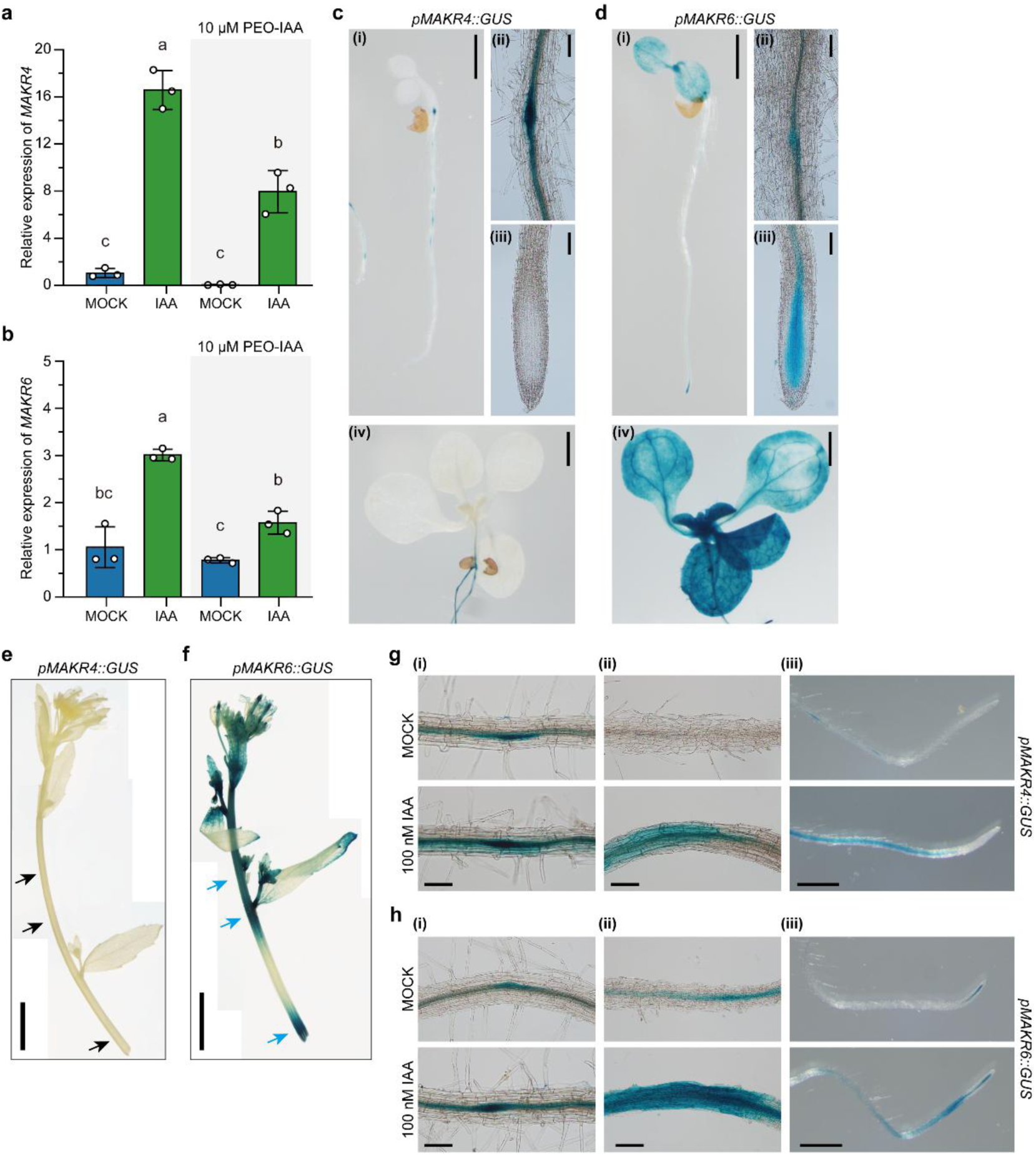
Expression pattern of *MAKR4* and *MAKR6* in plants. **a, b** qRT-PCR analyses of *MAKR4* and *MAKR6* transcription under treatments. 5-day-old seedlings were incubated for 4 hours with 1 μM IAA, MOCK, or 10 μM PEO-IAA, either alone or in combination, before RNA extraction. Relative expression levels of *MAKR4* and *MAKR6* were normalized to MOCK controls. **c-f** Expression patterns of *MAKR4* and *MAKR6* in plants revealed by GUS staining. *pMAKR4::GUS* (c, e) and *pMAKR6::GUS* (d, f) were analyzed in 4-day-old plants (c-i, c-ii, c-iii, d-i, d-ii, and d-iii), 10-day-old plants (c-iv and d-iv), and young shoots (e-f). *MAKR4* shows no obvious expression (black arrows), whereas *MAKR6* is strongly expressed in stems (blue arrows). **g, h** Auxin effects on *MAKR4* and *MAKR6* expression. *pMAKR4::GUS* (g) and *pMAKR6::GUS* (h) seedlings (4-day-old) were treated with auxin before GUS staining. Data represent results from at least three independent biological replicates, with consistent *GUS* expression patterns observed across >15 seedlings per line (n>3). Different letters indicate statistical differences (a, b, two-way ANOVA with Tukey’s HSD, P<0.05). Exact P values for all comparisons are provided in Source Data 1. Scale bars: 100 μm (c-ii, c-iii, d-ii, d-iii, g-i, g-ii, h-i, and h-ii), 2 mm (c-i, c-iv, d-i, d-iv, g-iii, and h-iii), and 50 mm (e, f).

**Extended Data Fig. 2.**
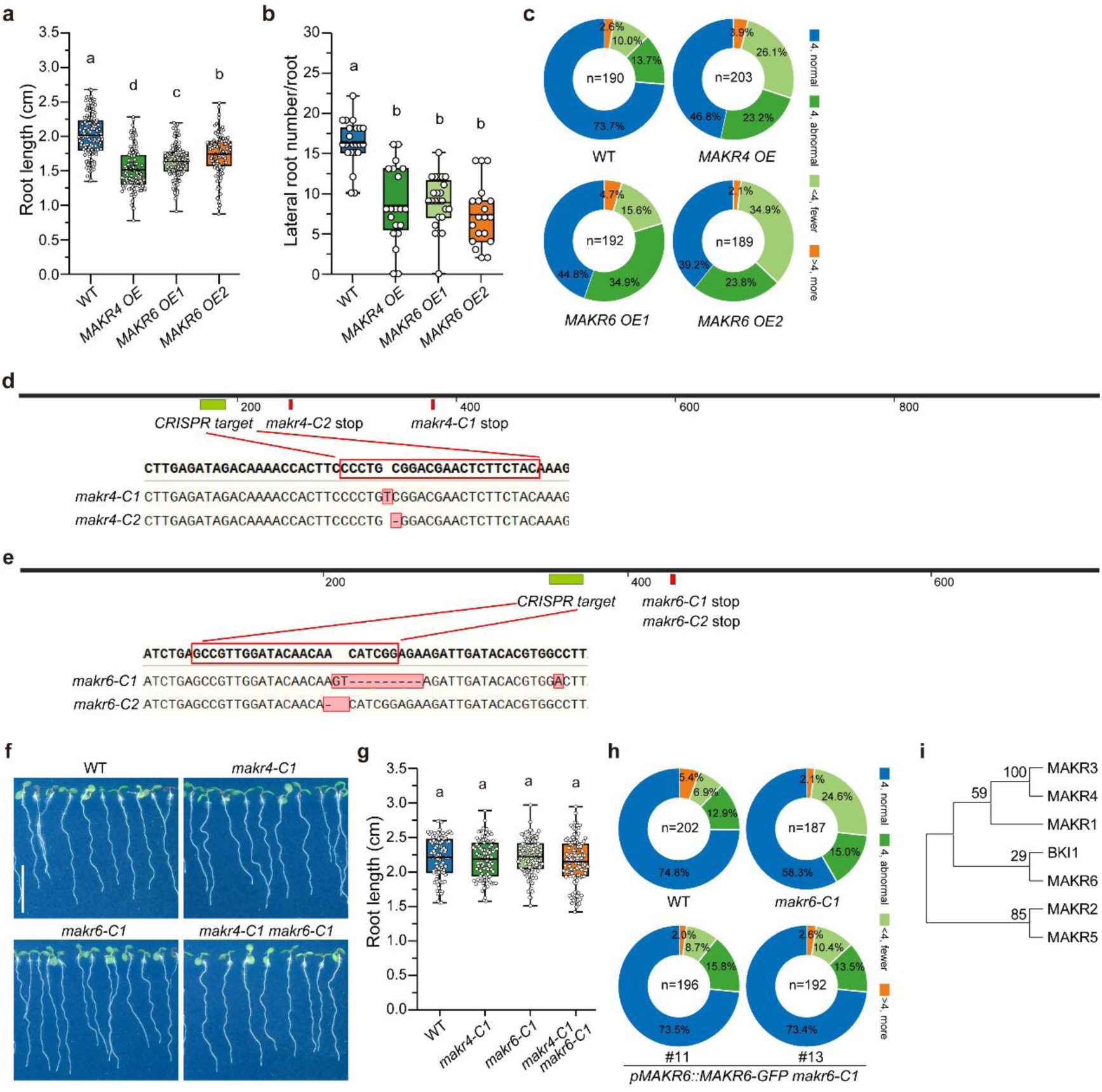
Distinct roles of MAKR4 and MAKR6 in plants. **a-c** Phenotypic analyses of *MAKR4* and *MAKR6* overexpression lines. Primary root length in 6-day-old seedlings (a), total lateral root number (b), and leaf vein patterns in 10-day-old seedlings (c) were analyzed in *p35S::MAKR4-GFP* (*MAKR4 OE*) and *p35S::MAKR6-GFP* (*MAKR6 OE1* and *OE2*). **d, e** CRISPR/Cas9-generated *makr4* and *makr6* mutants. Upper panels: gene map (coding sequence); lower panels: sequencing results of PCR-amplified fragments from mutant plants. **f, g** Root growth of *makr* mutants. 6-day-old seedlings were scanned for analysis. **h** Native expression of *MAKR6-GFP* rescued the defective leaf venation pattern in 10-day-old *makr6*. Sample sizes are indicated in the panel. **i** The phylogenetic tree shows that MAKR4 and MAKR6 cluster separately. Full-length protein sequences were aligned, and bootstrap values are indicated. Box plots show the interquartile range (boxes), the mean (central line), and the min-max range (whiskers), with open circles representing individual measurements. Sample sizes: n≥120 (a), n≥18 (b), and n≥85 (g). Abbreviations: OE, overexpression. Different letters indicate statistical differences (a, b, g, one-way ANOVA with Tukey’s HSD, P<0.05). Exact P values for all comparisons are provided in Source Data 1. Scale bar: 1 cm (f).

**Extended Data Fig. 3.**
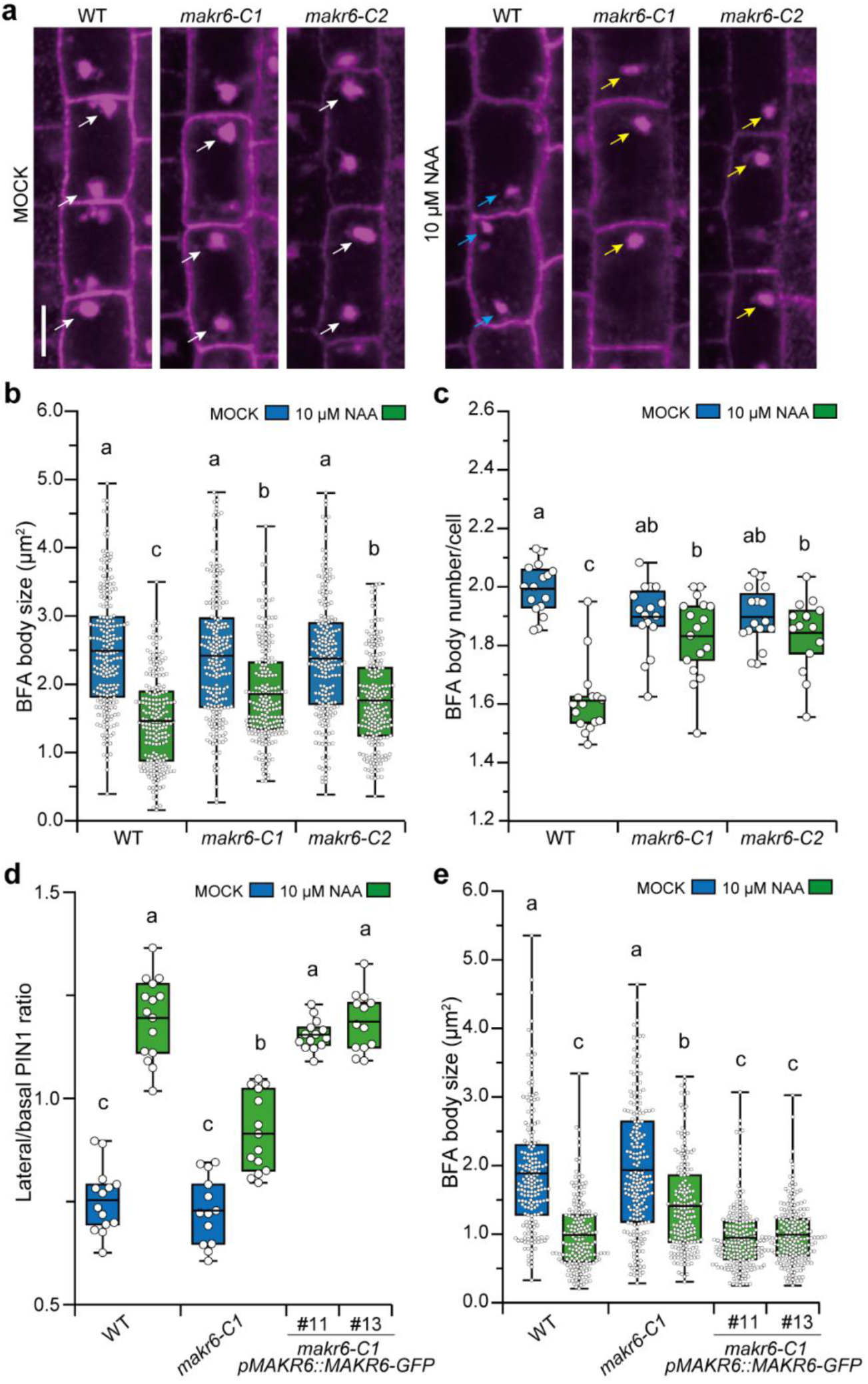
Complementation of auxin-sensitive PIN1 trafficking and repolarization by *pMAKR6::MAKR6-GFP*. **a-c** Representative images (a), quantification of individual BFA body size (b), and number of BFA bodies per cell (c) in pericycle cells are shown. White arrows show BFA bodies in typical size; blue arrows indicate smaller ones under auxin treatment; yellow arrows mark the abnormal BFA bodies. **d, e** Rescue of PIN1 lateralization (d) and BFA body size (e) in 6-day-old *pMAKR6::MAKR6-GFP* plants. PIN1 lateralization was calculated as the ratio of the inner lateral signal to the rootward basal signal. Box plots show the interquartile range (boxes), mean (central lines), and min-max range (whiskers). Open circles represent individual measurements from single BFA bodies (b, e) or averaged values per root (c, d). Data were collected from >150 cells across ≥10 roots per genotype, from one biological replicate of ≥2 biological replicates. Different letters indicate statistical differences (b, c, two-way ANOVA; d, e, one-way ANOVA with Tukey’s HSD, P<0.05). Exact P values for all comparisons are provided in Source Data 1. Scale bar: 5 μm (a).

**Extended Data Fig. 4.**
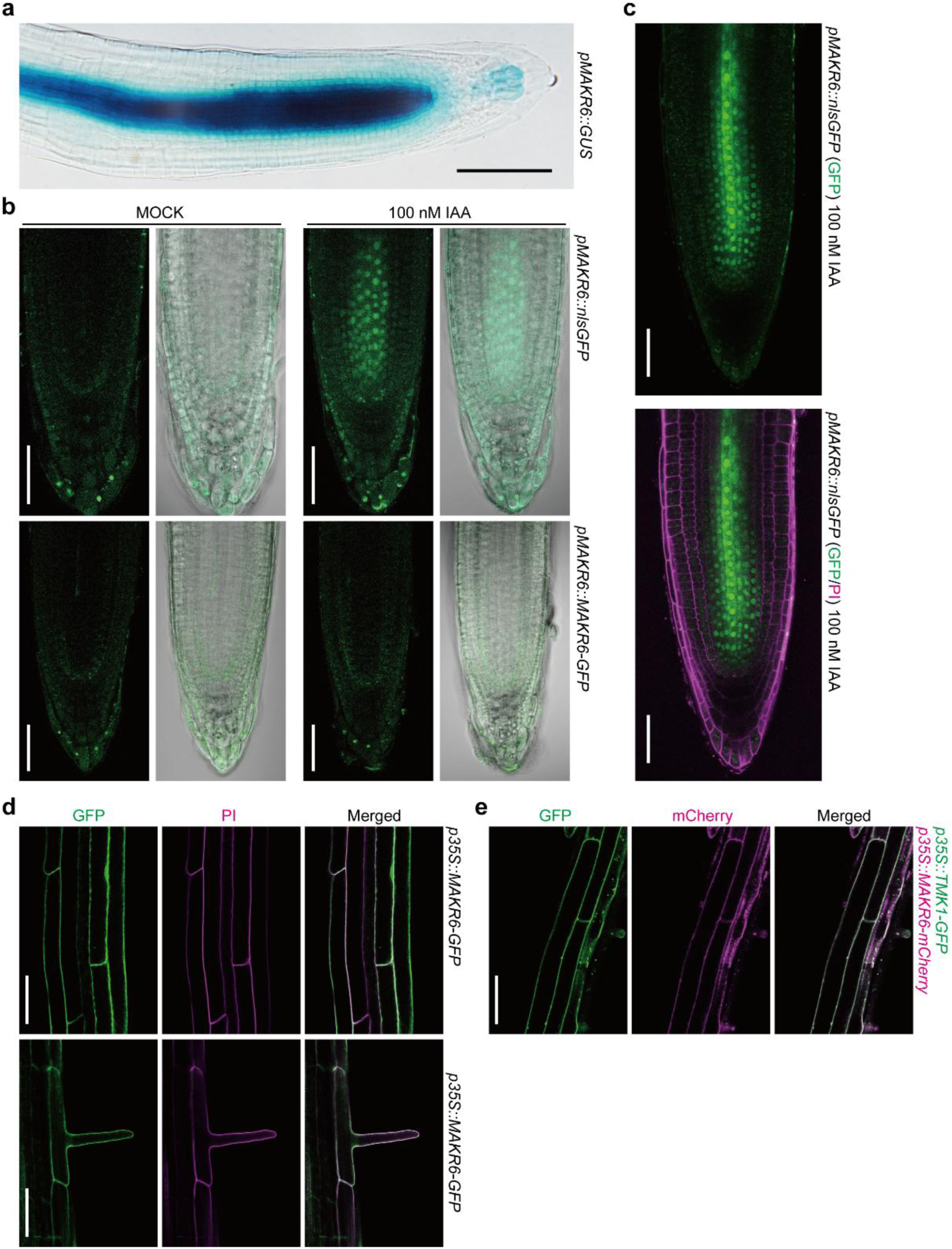
*MAKR6* expression pattern in root tips and MAKR6 subcellular localization at the plasma membrane. **a** Histochemical staining of 5-day-old *pMAKR6::GUS* showing *MAKR6* expression in root tips. **b, c** Representative confocal images of *pMAKR6::nlsGFP* and *pMAKR6::MAKR6-GFP* 5-day-old seedlings were treated with 100 nM IAA or MOCK solution for 4 hours before imaging. **d, e** Plasma membrane localization of MAKR6 in epidermal cells and root hair cells. 5-day-old seedlings of *p35S::MAKR6-GFP* or *p35S::TMK1-GFP;p35S::MAKR6-mCherry* was used, respectively. Consistent results were observed in at least 15 plants from each independent line (n>3). Abbreviations: PI, propidium iodide. Scale bars: 100 μm (a) and 50 μm (b-e).

**Extended Data Fig. 5.**
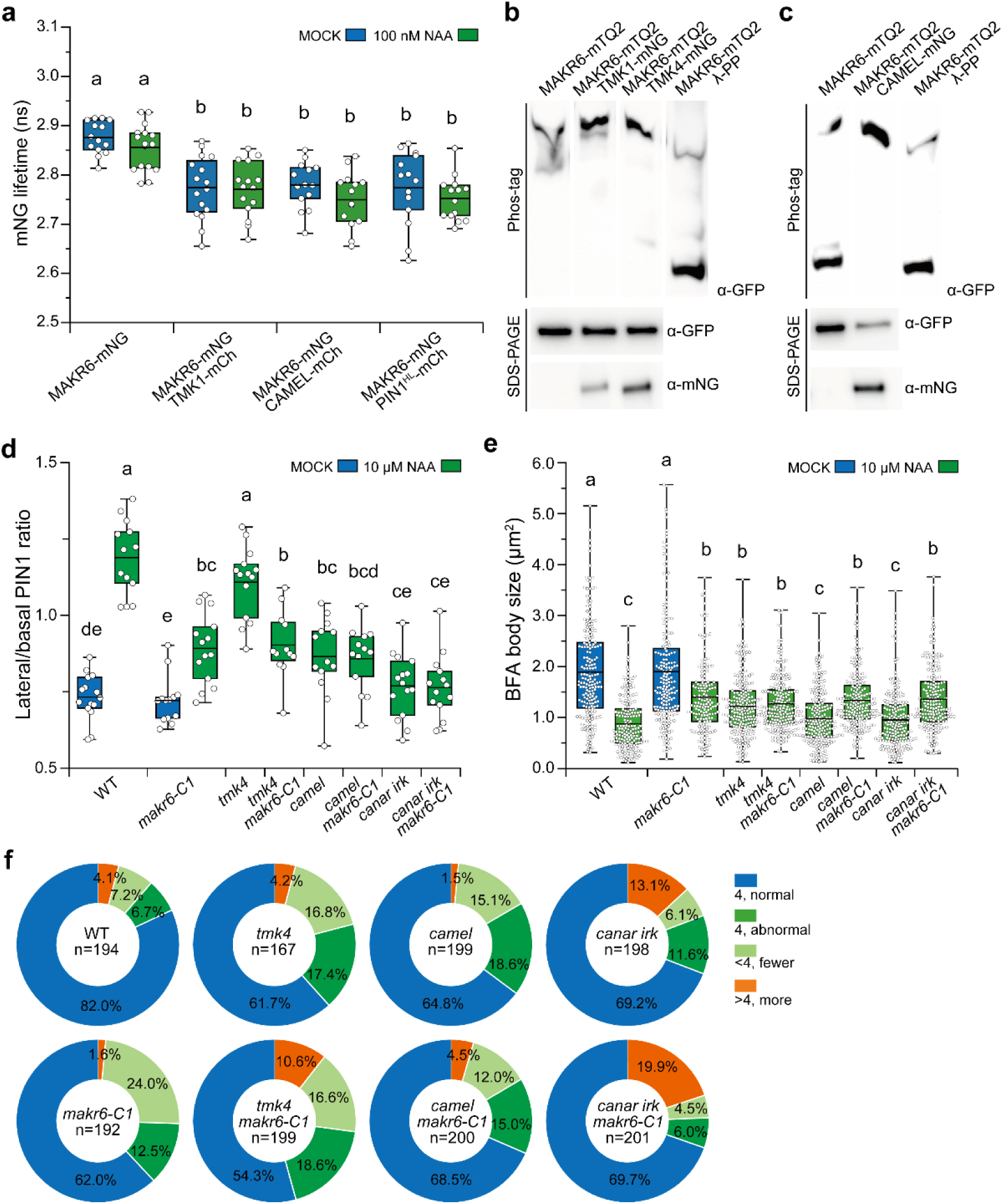
MAKR6 phosphorylation and genetic interactions with TMK, CAMEL, and CANAR receptors. **a** FLIM analysis showing no detectable auxin effect on MAKR6 interactions with TMK1, CAMEL, and PIN1 in protoplasts. NAA or MOCK solution was applied 2 hours before imaging. **b, c** TMK- and CAMEL-mediated phosphorylation of MAKR6 in Phos-tag analysis. MAKR6-mTQ2 alone or in combination with TMK/CAMEL was co-expressed in protoplasts. The upper and lower panels show Phos-tag gels and corresponding SDS-PAGE immunoblots, respectively. Uncropped blots are provided in Source Data 2. **d-f** Genetic interactions of MAKR6 with TMK, CAMEL, and CANAR receptors in regulating auxin-sensitive PIN1 repolarization, trafficking, and vein patterning. PIN1 immunostaining (d, e) was performed in 4-day-old seedlings and cotyledon patterns (f) in 10-day-old seedlings. Box plots show the interquartile range (boxes), mean (central lines), and min-max range (whiskers). Data were collected from ≥14 cells (a), 12 roots (d), and 160 BFA bodies or the indicated sample size (f), from one representative biological replicate of at least two independent biological replicates. Abbreviations: mCh, mCherry; mNG, mNeonGreen; mTQ2, mTurquoise2; λ protein phosphatase (λ-PP). Different letters indicate statistical differences (a, two-way ANOVA; d, e, one-way ANOVA with Tukey’s HSD, P<0.05). Exact P values for all comparisons are provided in Source Data 1.

**Extended Data Fig. 6.**
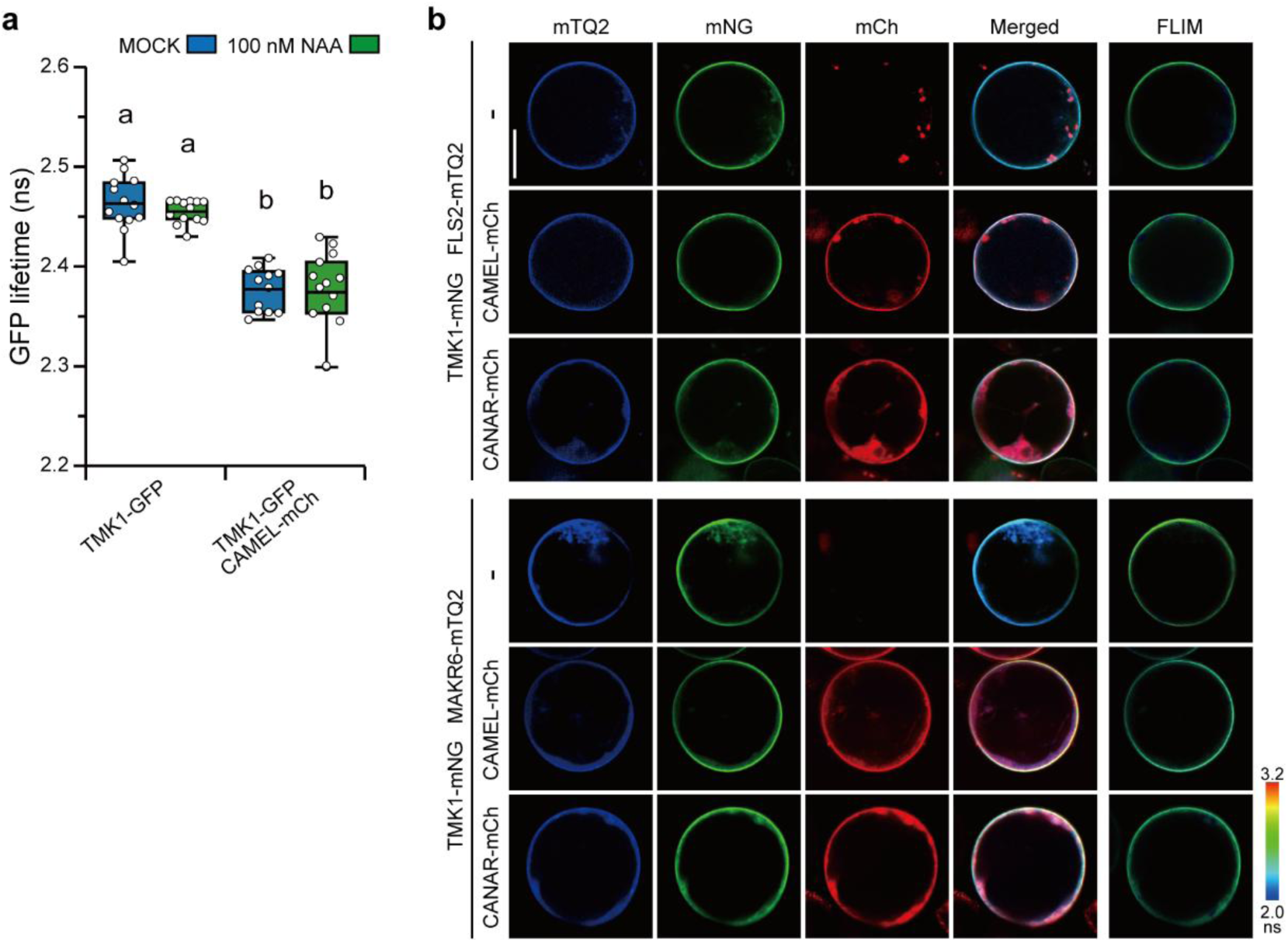
No obvious modulation of auxin and MAKR6 on the interaction between TMK1 and CAMEL-CANAR. **a** FLIM quantification showing no obvious auxin effect on the interaction between TMK1 and CAMEL. NAA or MOCK solution was applied to protoplasts expressing the indicated proteins 2 hours before imaging. **b** Representative FLIM images in Arabidopsis protoplasts transiently expressing TMK1-mNG either alone or together with CAMEL/CANAR-mCh, in the presence of MAKR6-mTQ2 or FLS2-mTQ2. Quantitative results are shown in Fig. 6g. Box plots show lifetime values, with boxes representing the interquartile range, central lines the mean, and whiskers the min-max range. Data were collected from ≥12 cells per group in a single representative biological replicate from at least 2 independent experiments. Abbreviations: mCh, mCherry; mNG, mNeonGreen; mTQ2, mTurquoise2. Different letters indicate statistical differences (a, two-way ANOVA with Tukey’s HSD, P<0.05). Exact P values for all comparisons are provided in Source Data 1. Scale bar, 20 μm. The colour scales indicate the mNG lifetime.

**Extended Data Fig. 7.**
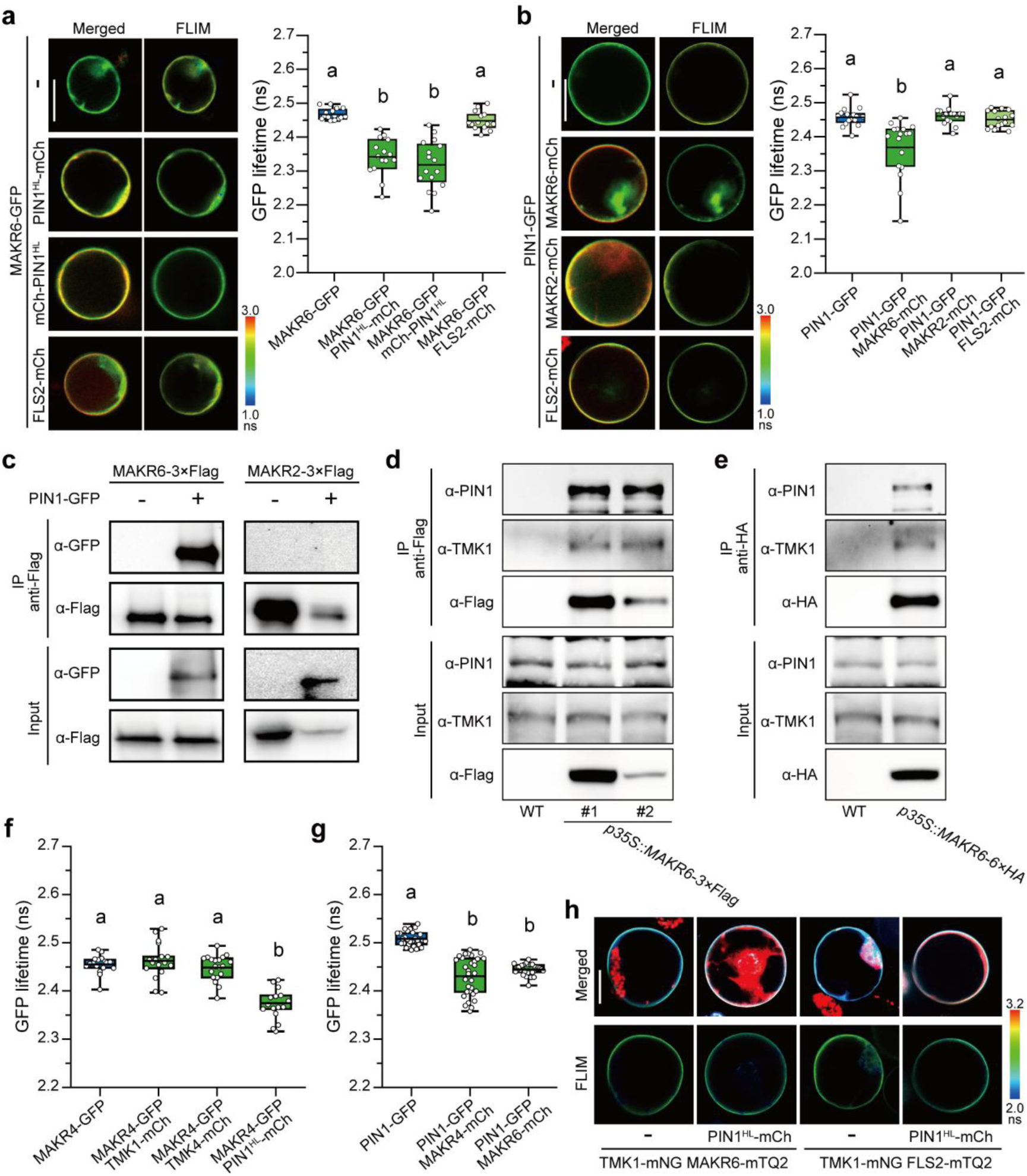
Interaction between MAKR4/6, TMK, and PIN1. **a, b** MAKR6 specifically interacts with PIN1 in FRET/FLIM analyses. Arabidopsis root protoplasts were co-transformed with *p35S::MAKR6-GFP*, *p35S::PIN1^HL^-mCh,* and *p35S::mCh-mPIN1^HL^* in combinations (a), or combined *p35S::PIN1-GFP*, *p35S::MAKR6-mCh,* and *p35S::MAKR2-mCh* (b). *p35S::FLS2-mCh* was used as a control. **c-e** Co-IP assays revealing the interaction of MAKR6 with TMK1 and PIN1. Immunoprecipitation was performed with anti-Flag beads using transient expression in tobacco leaves (c), anti-Flag or anti-HA beads using Arabidopsis seedlings expressing *p35S::MAKR6-3×Flag* and *p35S::MAKR6-6×HA* (d, e). **f, g** FLIM quantification showing that MAKR4 does not significantly interact with TMK1/4, whereas it does interact with PIN1, similar to MAKR6. **h** Representative images in the FRET/FLIM assay testing the MAKR6 effect on the interaction between TMK1 and PIN1. Quantitative results are shown in Fig. 6h. Box plots show GFP lifetime, with boxes representing the interquartile range, central lines the mean, and whiskers the min-max range. Open circles denote individual cell measurements (n≥13 cells per group). Abbreviations: mCh, mCherry; mNG, mNeonGreen; mTQ2, mTurquoise2. Different letters indicate statistical differences (a, b, f, g, one-way ANOVA with Tukey’s HSD, P<0.05). Exact P values for all comparisons are provided in Source Data 1. Scale bar: 20 µm. Colour scales indicate GFP lifetime (a, b) and mNG lifetime (h).

**Extended Data Fig. 8.**
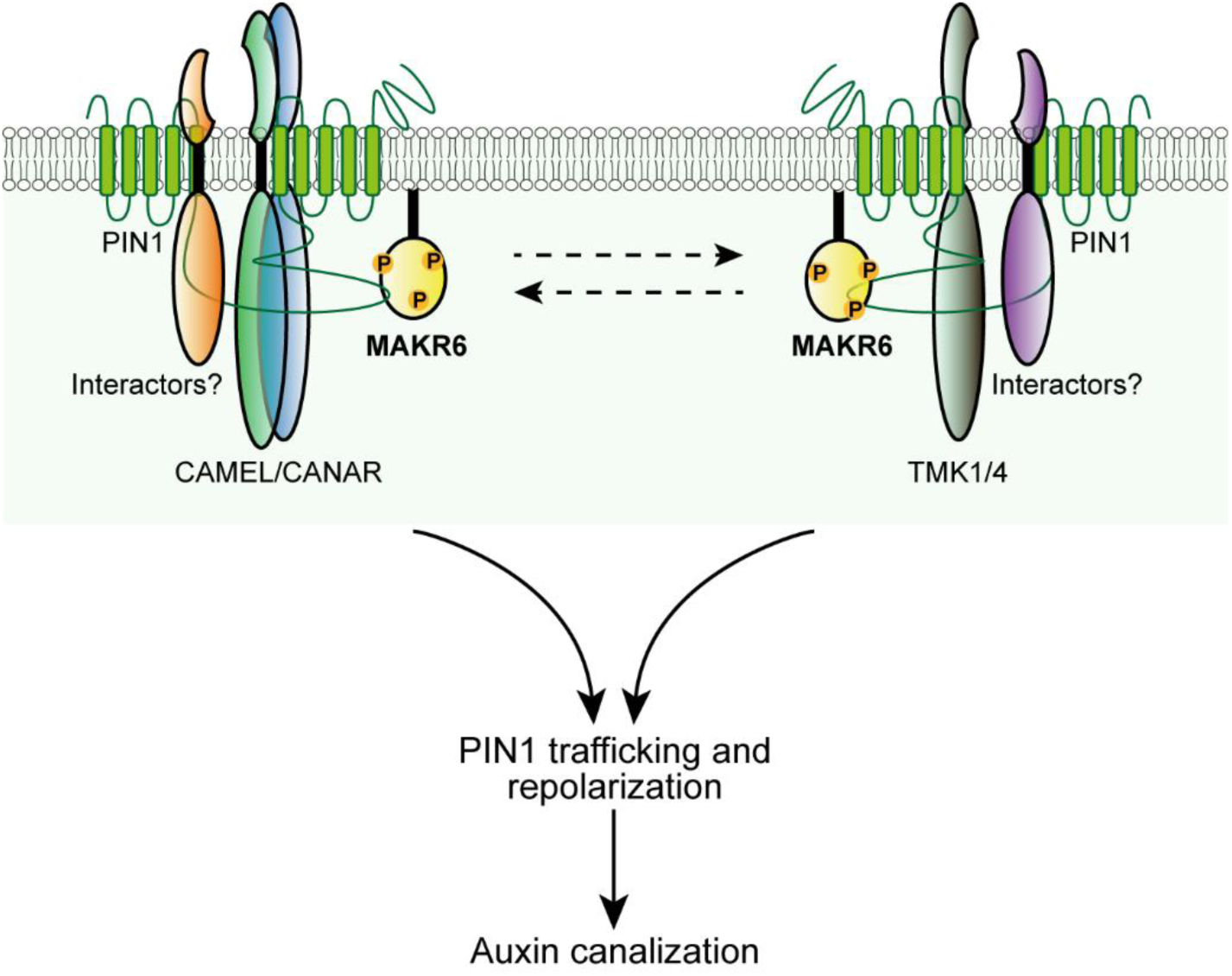
Working model for MAKR6 function in auxin canalization. The diagram illustrates a proposed role for MAKR6 in coordinating distinct receptor-like kinase complexes to regulate auxin canalization. MAKR6 interacts with the CAMEL/CANAR and TMK1/4 kinase complexes, as well as with PIN1, likely within dynamic or distinct interaction complexes at the plasma membrane. These pathways converge on the regulation of PIN1 dynamics, thereby promoting auxin canalization-dependent processes, including venation and vascular regeneration. TMK- and CAMEL-associated signalling pathways regulate MAKR6 through phosphorylation, potentially downstream of ligand perception and pathway activation. Dashed arrows indicate potential cross-regulation between receptor complexes, and question marks denote additional components, such as co-receptors, that remain to be identified.

## Supplementary information

**Supplementary Table 1.** Constructs generated in this work.

|  |  |
| --- | --- |
| *gMAKR4*/pDONR221 | Entry clone, *MAKR4* genomic (no stop codon) |
| *gMAKR6*/pDONR221 | Entry clone, *MAKR6* genomic (no stop codon) |
| *pMAKR4*/pDONRP4P1r | Entry clone, *MAKR4* promoter |
| *pMAKR6*/D-Topo | Entry clone, *MAKR6* promoter |
| *pMAKR6*/pGGA000 | Entry clone, *MAKR6* promoter |
| *gMAKR6*/pGGC000 | Entry clone, *MAKR6* genomic (no stop codon) |
| *gMAKR4*/pB7FWG2 | Expression clone for *MAKR4* overexpression |
| *gMAKR6*/pB7FWG2 | Expression clone for *MAKR6* overexpression |
| *pMAKR4::GUS*/pK7m24GW | Expression clone for *pMAKR4::GUS* |
| *pMAKR6*/pB7GUSWG0 | Expression clone for *pMAKR6::GUS* |
| *pMAKR6-MAKR6-GFP*/pBLAX | Expression clone for *pMAKR6-MAKR6-GFP* |
| *pMAKR6*/pGWB465 | Expression clone for *pMAKR6::nlsGFP* |
| *CAMEL*/pB7HAWG2 | Expression clone for CAMEL-6×HA in Co-IP |
| *CANAR*/pB7HAWG2 | Expression clone for CANAR-6×HA in Co-IP |
| *gTMK1*/pB7HAWG2 | Expression clone for TMK1-6×HA in Co-IP |
| *gTMK4*/pB7HAWG2 | Expression clone for TMK4-6×HA in Co-IP |
| *gMAKR6*/pB7HAWG2 | Expression clone for MAKR6-6×HA in Co-IP |
| *CAMEL*/pB7FLAGWG2 | Expression clone for CAMEL-3×Flag in Co-IP |
| *CANAR*/pB7FLAGWG2 | Expression clone for CANAR-3×Flag in Co-IP |
| *gTMK1*/pB7FLAGWG2 | Expression clone for TMK1-3×Flag in Co-IP |
| *gTMK4*/pB7FLAGWG2 | Expression clone for TMK4-3×Flag in Co-IP |
| *MAKR2*/pB7FLAGWG2 | Expression clone for MAKR2-3×Flag in Co-IP |
| *gMAKR6*/pB7FLAGWG2 | Expression clone for MAKR6-3×Flag in Co-IP |
| *gMAKR6*/pCoE2GWmNG7 | Expression clone for MAKR6-mNG in FRET/FLIM |
| *gTMK1*/pCoE2GWmCh7 | Expression clone for TMK1-mCh in FRET/FLIM |
| *gTMK4*/pCoE2GWmCh7 | Expression clone for TMK4-mCh in FRET/FLIM |
| *CAMEL*/pCoE2GWmCh7 | Expression clone for CAMEL-mCh in FRET/FLIM |
| *CANAR*/pCoE2GWmCh7 | Expression clone for CANAR-mCh in FRET/FLIM |
| *FLS2*/pCoE2GWmCh7 | Expression clone for FLS2-mCh in FRET/FLIM |
| *CANAR*/pB7FWG2 | Expression clone for CANAR-GFP in FRET/FLIM |
| *CAMEL*/pB7FWG2 | Expression clone for CAMEL-GFP in FRET/FLIM |
| *gTMK1*/pB7FWG2 | Expression clone for *TMK1* overexpression |
| *gTMK1*/pB7MWG2 | Expression clone for TMK1-mCh in FRET/FLIM |
| *gTMK4*/pB7MWG2 | Expression clone for TMK4-mCh in FRET/FLIM |
| *CAMEL*/pB7MWG2 | Expression clone for CAMEL-mCh in FRET/FLIM |
| *FLS2*/pB7MWG2 | Expression clone for FLS2-mCh in FRET/FLIM |
| *MAKR2*/pB7MWG2 | Expression clone for MAKR2-mCh in FRET/FLIM |
| *MAKR4*/pB7MWG2 | Expression clone for MAKR4-mCh in FRET/FLIM |
| *gMAKR6*/pB7MWG2 | Expression clone for MAKR6-mCh in FRET/FLIM and *MAKR6* overexpression |
| *gMAKR6*/pCoE2GWmTQ27 | Expression clone for MAKR6-mTQ2 in FRET/FLIM |
| *gMAKR6*/pCoE2GWmTQ27-<br>*gTMK1*/2GWmNG7 | Expression clone for MAKR6-mTQ2 and TMK1-mNG in FRET/FLIM |
| *gMAKR6*/pCoE2GWmTQ27-<br>*gTMK4*/2GWmNG7 | Expression clone for MAKR6-mTQ2 and TMK4-mNG in FRET/FLIM |
| *gMAKR6*/pCoE2GWmTQ27-<br>*CAMEL*/2GWmNG7 | Expression clone for MAKR6-mTQ2 and CAMEL-mNG in FRET/FLIM |
| *FLS2*/pCoE2GWmTQ27-<br>*gTMK1*/2GWmNG7 | Expression clone for FLS2-mTQ2 and TMK1-mNG in FRET/FLIM |

